# Weakening of subcortical and strengthening of cortical visual pathways across early adolescence

**DOI:** 10.1101/2025.09.05.674387

**Authors:** Arshiya Sangchooli, Elise G. Rowe, Robert E. Smith, Iroise Dumontheil, Marta I. Garrido

## Abstract

**Background:** Mounting evidence suggests that amygdalar nuclei receive visual information via both a well-characterized cortical pathway through the inferior temporal cortex and a subcortical route through the superior colliculus and pulvinar. This subcortical pathway may facilitate rapid responses to salient visual stimuli and could explain phenomena such as blindsight. However, controversies remain about the organization of the subcortical pathway, its role in visual processing, and how the cortical and subcortical pathways mature across development.

**Methods:** To address these questions we used longitudinal diffusion magnetic resonance imaging (dMRI) data from 4361 participants in the Adolescent Brain Cognitive Development (ABCD) study, reconstructing every major segment of the cortical and subcortical amygdala pathways. We tested the existence of the subcortical pathway against null tractography models, characterized cortical and subcortical pathways development across early adolescence, and investigated their association with visual processing speed.

**Results:** We provide evidence for the existence of bilateral pulvinar-amygdala pathways against a null model (all _*p*_ < 0.001, corrected). While cortical tracts involving the primary and extrastriate visual cortex and the inferior-temporal cortex strengthened with chronological age and over pubertal development, we demonstrate that subcortical pulvinar-amygdala connectivity decreased over pubertal development. Greater connectivity strength of the right pulvinar-amygdala tract was associated with faster responses on a visual task for both emotional face and place stimuli, a relationship also seen for cortical tracts.

**Conclusion:** This study provides evidence for the existence of pulvinar to amygdala tracts in the largest sample of adolescent participants studied to date. Greater connectivity in both cortical and subcortical tracts were associated with faster reaction time on a visual task, but further work will be needed to investigate the specificity of this association in terms of both task and tract. In line with the hypothesized importance of the subcortical pathway in early development, we show that the developmental trajectories of cortical and subcortical pathways diverge and highlight the influence of pubertal development, with cortical pathways generally strengthening and subcortical pathways weakening across early adolescence.

## Introduction

Amygdalar nuclei play a wide array of roles in processing sensory information across humans and many other non-human species. The amygdala is known to both receive and gate primary and secondary sensory inputs, integrate these with higher-cortical information to process the valence of the stimuli, and use such information to bias attention and rapidly shape behavioral responses (Gothard 2020; Janak and Tye 2015). Amygdalar processing is crucial in reward evaluation, conditioning, and social processing, such as processing facial expressions (Fast and McGann 2017; Gothard and Fuglevand 2022); though the most intensively investigated amygdalar function is arguably learning about and responding to threat (Li 2019).

In primates, the amygdalae receive emotionally salient visual information – such as threatening scenes or face stimuli – from the ventral stream: from the retina this information flows to the lateral geniculate nucleus (LGN) and then the primary and extrastriate visual cortices, finally reaching the amygdala via its rich cortical connections with the inferior temporal cortex (Stefanacci and Amaral 2002; Ungerleider, Gaffan, and Pelak 1989). In addition, there is accumulating evidence for a rapid subcortical pathway for visual stimuli, primarily through the superior colliculus and pulvinar, in rodents, nonhuman primates, and humans (McFadyen, Mattingley, and Garrido 2019). This so-called “low road” pathway has garnered increased attention since the discovery of subcortical tracts for threatening stimuli to the amygdala in rodents that bypass the primary auditory cortex (LeDoux 1998). The subcortical pathway could explain amygdalar responses to emotionally salient visual stimuli in patients with damaged primary visual cortices (Burra et al. 2013) but residual vision (or “blindsight”) (McFadyen, Dolan, and Garrido 2020). This pathway may also account for the extremely fast amygdalar responses to fearful facial expressions (Méndez-Bértolo et al. 2016), and early neonatal responses to threatening visual stimuli (Ball and Tronick 1971) and to faces (Johnson, 2005). Major tracts in the proposed subcortical visual pathway, along with the canonical cortical pathway, are presented in *Figure 1*. It is thought that the subcortical pathway receives direct retinal input to the superior colliculi (SC) and pulvinar (McFadyen, Dolan, and Garrido 2020), though there is also some evidence for LGN-SC connections (Basso & May, 2017) across several species (Brauer & Schober, 1982; Monavarfeshani et al., 2017; Nakamura & Itoh, 2004).

**Figure 1:**
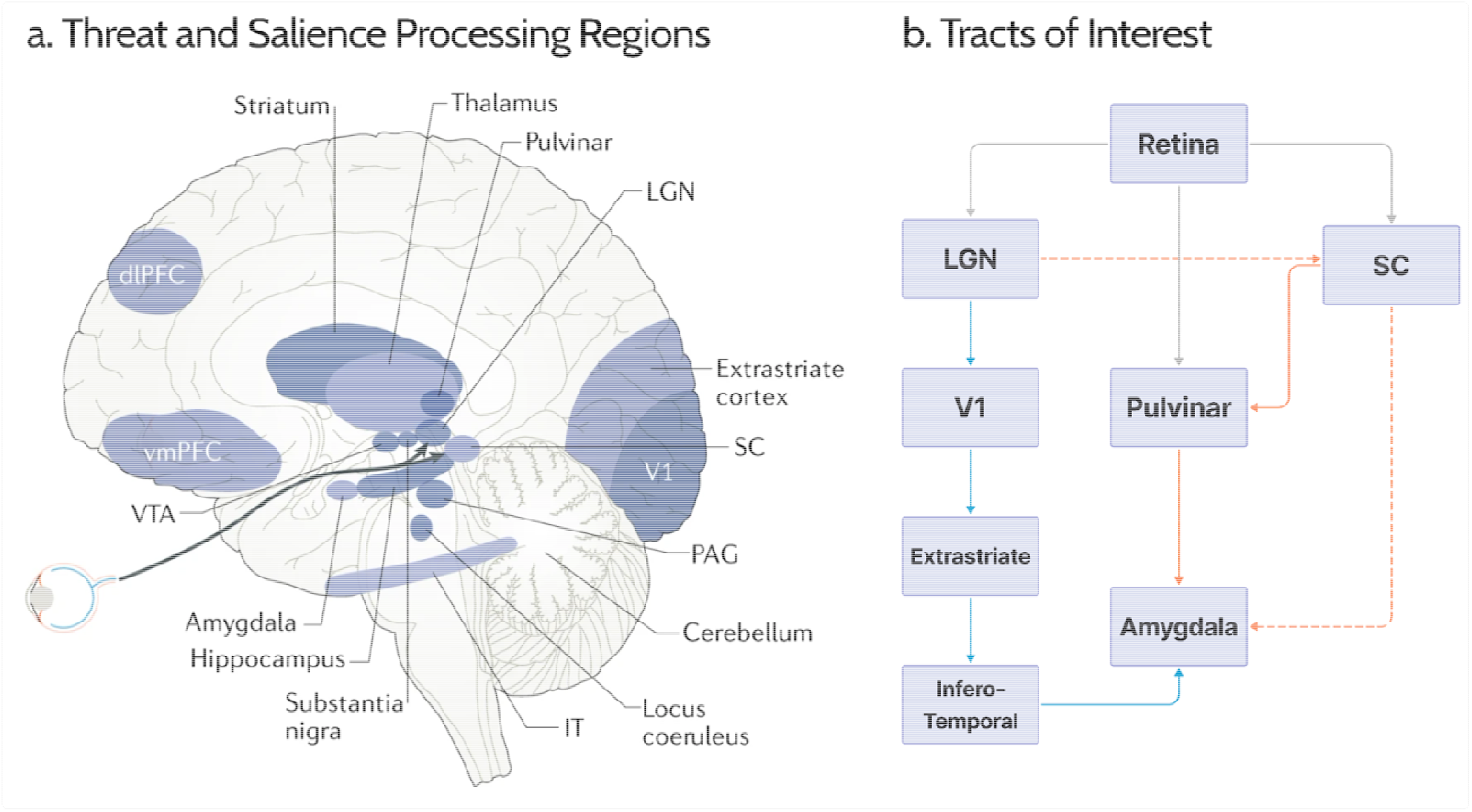
The circuitry of salient visual processing. a) Regions involved in processing and responding to threatening or otherwise salient visual information. b) Schematic of the human cortical and subcortical visual pathways to the amygdala. Major visual tracts in the cortical (blue) and subcortical (orange) pathways to the amygdala. Dashed lines represent tracts with less support in the literature. Retinal tracts (in gray) were not investigated due to the difficulty of tracing tracts from the retina. SC: superior colliculus; LGN: lateral geniculate nucleus; dlPFC: dorsolateral prefrontal cortex: ES, extrastriate cortex; IT: inferotemporal cortex; PAG: periaqueductal gray; V1: primary visual cortex; vmPFC: ventromedial prefrontal cortex; VTA: ventral tegmental area. Figure adapted with permission from Springer Nature, License Number: 5796460438365; from McFadyen et al. (2020).

Given most of the growing literature on the subcortical pathway has been focused on providing evidence for its *existence and functional role* (Elorette et al. 2018; McFadyen, Mattingley, and Garrido 2019), the *development* of these tracts remains under-investigated. This is despite abundant evidence that the human amygdala matures rapidly throughout adolescence (Zhou et al. 2021), possibly because the amygdala has an exceptionally high density of steroid hormone receptors, and is therefore sensitive to pubertal development (Goddings et al. 2019). A few recent diffusion magnetic resonance imaging (dMRI) studies have suggested that amygdalar white matter and major amygdalar connections to cortical and diencephalic regions mature substantially during adolescence, with myelination and structural connectivity indices generally showing a strengthening of major white matter bundles in this period (Azad et al. 2021; Goetschius et al. 2019). However, the development of the subcortical visual pathway to the amygdala remains unexplored and, importantly, there is reason to believe that the development trajectories of visual pulvinar and cortical tracts may differ. For instance, pulvinar tracts seem to provide an important shortcut for visual information at least early in life, playing an important role in the initial development of the primate dorsal stream. During primate development, these tracts prune as the geniculostriate pathway matures, unless the latter is damaged (Warner et al. 2015; Bridge, Leopold, and Bourne 2016), in which case the pulvinar tract remains unpruned in what appears to be a mechanism of compensatory redundancy for residual vision (McFadyen, Dolan, Garrido, 2021). While understanding the development of these pathways could provide critical insights into an understudied aspect of subcortical brain development, there have been no studies to date on the developmental trajectory of subcortical visual tracts to the amygdala in humans.

In this study, we investigated multiple hypotheses relating to amygdala pathways in *N*=4361 young adolescents from the longitudinal Adolescent Brain Cognitive Development (ABCD) study. Firstly, we investigated evidence for the existence of different subcortical amygdala tracts by comparing their dMRI tractography reconstruction to corresponding null tractography (Morris et al. 2008). Secondly, we asked whether the subcortical visual pathway enables rapid relaying of salient visual information to the amygdala (McFadyen, Mattingley, and Garrido 2019), by testing a putative association between structural connectivity of subcortical tracts and the speed of emotional face processing. Finally, we investigated the developmental trajectories of quantitative structural connectivity in cortical and subcortical visual pathways, considering the influences of both pubertal development and chronological age.

## Methods

### Participants

This study used data from the ABCD study, one of the largest longitudinal studies of brain development (Casey et al. 2018), with data collected at multiple sites in the US and using three different scanners with consistent imaging and behavioral data collection protocols across sites. Participants were 9-10 years old at baseline and 11-12 at the second imaging time point. Participants with quality-checked diffusion MRI, and T1-weighted and T2-weighted images at both time points who did not meet any exclusion criteria were included in the analyses (N=4361). Exclusion criteria were: 1) history of traumatic brain injury with loss of consciousness; 2) epilepsy, seizure, cerebral palsy, multiple sclerosis, or any other major neurological condition; 3) any visible variation or abnormality in structural brain scans; 4) high motion, defined as >5 mm on average during the diffusion scan as per previous studies in adolescents (Leong et al. 2021); 5) poor diffusion MRI coverage of the brain at either time point, defined as a dice coefficient below 0.95 between participant and template masks after registering and transforming derivative diffusion image data to a common template space; 6) poor coverage of either superior colliculus at either timepoint (<0.75% of either superior colliculus contained in a brainstem mask derived from Freesurfer; Iglesias et al. 2015). One participant was excluded due to very poor registration to the template. Demographic characteristics of included participants are presented in **Table 1**.

**Table 1:**
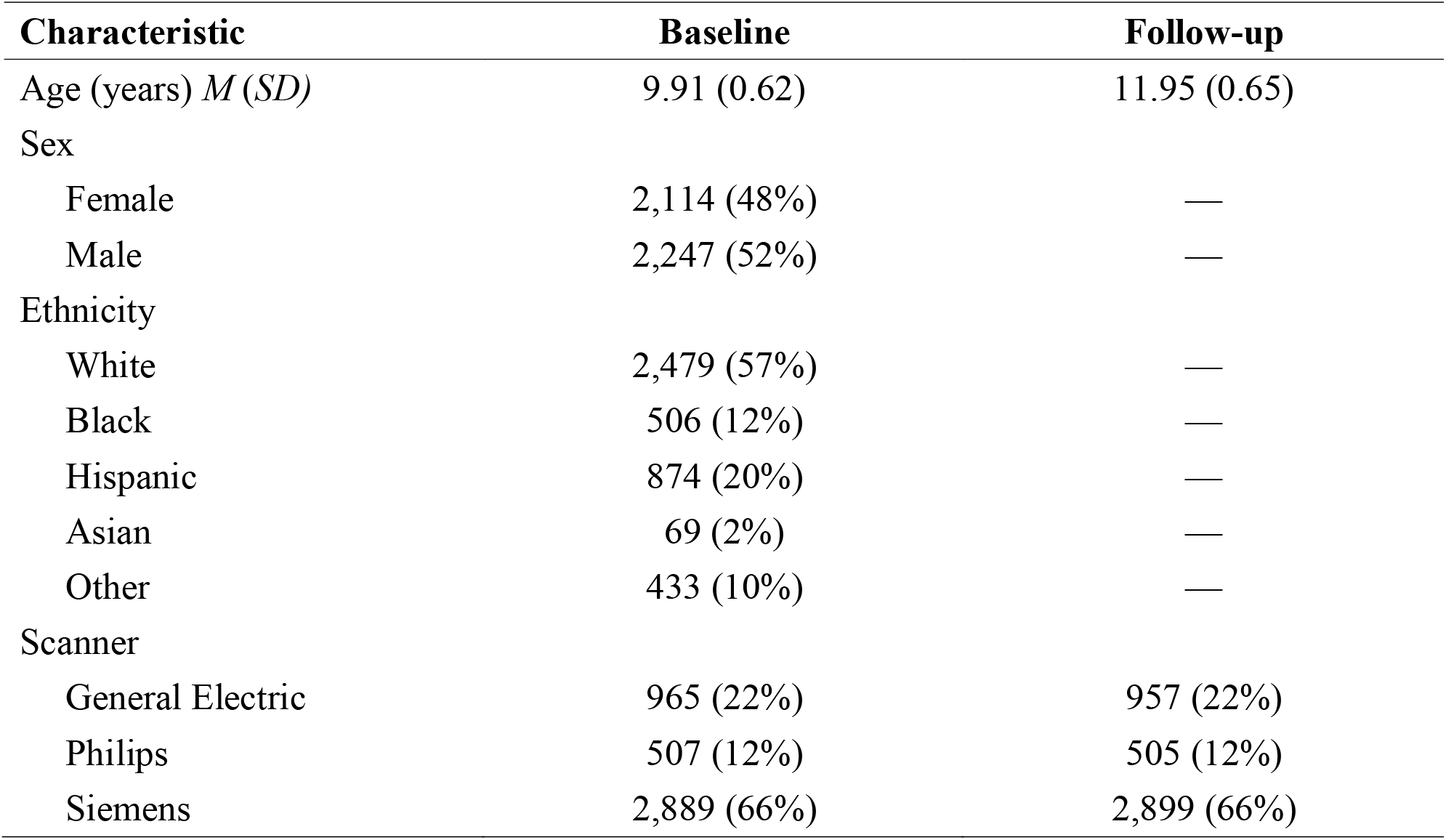
Participant demographics and scanner types at baseline and follow-up timepoints.

| Characteristic | Baseline | Follow-up |
| --- | --- | --- |
| Age (years) *M (SD)* | 9.91 (0.62) | 11.95 (0.65) |
| Sex |  |  |
| Female | 2,114 (48%) | — |
| Male | 2,247 (52%) | — |
| Ethnicity |  |  |
| White | 2,479 (57%) | — |
| Black | 506 (12%) | — |
| Hispanic | 874 (20%) | — |
| Asian | 69 (2%) | — |
| Other | 433 (10%) | — |
| Scanner |  |  |
| General Electric | 965 (22%) | 957 (22%) |
| Philips | 507 (12%) | 505 (12%) |
| Siemens | 2,889 (66%) | 2,899 (66%) |

### Structural MRI Processing

T1- and T2-weighted MRI images were obtained from the ABCD study. Details of the image acquisition and minimal preprocessing have been previously described (Hagler et al. 2019). Structural scans were automatically processed with the FreeSurfer image analysis suite, version 7.2.0 (Fischl 2012). The technical details of these procedures are described in prior publications, and include motion correction, removal of non-brain tissue, tissue-type segmentation, automated parcellation of subcortical and cortical structures, and cortical surface estimation, among other steps (Dale, Fischl, and Sereno 1999; Desikan et al. 2006; Fischl, Liu, and Dale 2001; Fischl, Salat, et al. 2004; Fischl, Sereno, and Dale 1999; Fischl et al. 1999; Fischl, van der Kouwe, et al. 2004; Segonne et al. 2004; Segonne, Pacheco, and Fischl 2007). We processed the structural scans obtained at the two time points from each participant with the longitudinal stream in FreeSurfer, which involves mapping the scans to an unbiased within-participant template space and initializing several processing steps with common information from the template to significantly increase reliability and statistical power (Reuter, Rosas, and Fischl 2010; Reuter et al. 2012). T2-weighted scans were provided in addition to T1-weighted images to improve the robustness of the pial surface estimation.

### Diffusion MRI Processing

Minimally processed multi-shell HARDI images registered to the structural scans were also obtained from the ABCD study, with acquisition and preprocessing details described by Hagler et al. (2019). Diffusion image data were further processed using a bespoke iterative approach that integrates B1 bias field correction, global intensity normalization, and brain mask derivation, exploiting information from multi-tissue decomposition of the diffusion signal (Jeurissen et al. 2014; Raffelt et al. 2017; Dhollander et al. 2021), with the brain mask derivation component performed using SynthStrip (Hoopes et al. 2022).

Diffusion MRI data were registered and transformed (without regridding) to the bias-field corrected and masked structural scans obtained with FreeSurfer, using a rigid-body boundary-based registration of mean *b*=0 images and the white matter segmentation obtained by FreeSurfer (Greve and Fischl 2009; Jenkinson et al. 2002; Jenkinson and Smith 2001) as implemented in FSL version 6.0.5.1 (Jenkinson et al. 2012). White matter, gray matter, and cerebrospinal fluid response functions were estimated for each individual using a data-driven approach (Dhollander et al. 2019); average response functions were calculated independently for each of the three scanner models given inconsistencies in acquisition parameters. Averaged response functions were used to estimate multi-tissue orientation distribution functions using all available *b*-value shells (Jeurissen et al. 2014; Tournier, Calamante, and Connelly 2007). Where not otherwise explicitly stated, analysis steps for diffusion MRI data were implemented using MRtrix3, version 3.0.3 (Tournier et al. 2019).

### Segmenting Regions of Interest

Segmentations of anatomical structures necessary for delineation of the bundles of interest were derived for each scanning session using image processing techniques tailored for each structure. (a) In the absence of individualized functional retinotopic mapping data, V1 and the extrastriate cortices (including V2, V3 and V4) were delineated by aligning flattened participant cortical surfaces to a retinotopy template (Wang et al. 2015) using FreeSurfer and the Neuropythy library (Benson 2022; Benson et al. 2014). This approach is suitable for early visual areas where there is approximate correspondence between anatomy and function such that probabilistic surface-based atlases perform satisfactorily (Alvarez, Parker, and Bridge 2019). (b) The pulvinar nuclei and LGN were segmented with a multi-spectral thalamic segmentation method using participant T1- and T2-weighted scans and a probabilistic atlas built with histological data, as implemented in FreeSurfer (Iglesias et al. 2018). (c) Amygdalar nuclei were segmented with another multi-spectral segmentation algorithm implemented in FreeSurfer (Saygin et al. 2017), which uses an atlas created from postmortem high-resolution structural scans. (d) The inferior temporal cortex was segmented by merging the relevant gyri (lateral occipito-temporal and inferior temporal) and sulci (anterior transverse collateral, lateral and medial occipito-temporal, and inferior temporal) from the Destrieux atlas in FreeSurfer (Destrieux et al. 2010), and taking their union with the VO1, VO2, PHC1 and PHC2 regions to ensure inclusion of these important visual areas of the inferior temporal cortex (see Wang et al. 2015 for precise anatomical locations). (e) The superior colliculi were delineated by non-linearly registering participant diffusion MRI data to the Illinois Institute of Technology (IIT) template and using a recently developed probabilistic atlas of brainstem nuclei (García-Gomar et al. 2019), available in the Brainstem Navigator toolkit (Bianciardi 2021). This atlas has been used in multiple recent brainstem tractography studies (García-Gomar et al. 2022; Singh et al. 2022).

### Tractography

Two different tractography experiments were performed: the first to assess the evidence for the existence of subcortical tracts, and the second to assess a) the relationship between tract structural connectivity and behavioral measures, and b) the development of structural connectivity in cortical and subcortical amygdala pathways. In both experiments, estimated white matter fibre orientation distributions were used to generate streamlines for each participant (at each timepoint) with the probabilistic iFOD2 algorithm (Tournier, Calamante, and Connelly 2010). We used default parameters, except maximum streamline length (set to 200 mm to allow for the reconstruction of the longest-range cortical connections) and minimum streamline length (set to 0.75 mm to allow reconstruction of short-range fibres). The major steps of the analysis pipeline are illustrated in **Figure 2** and described in detail in the following subsections.

**Figure 2:**
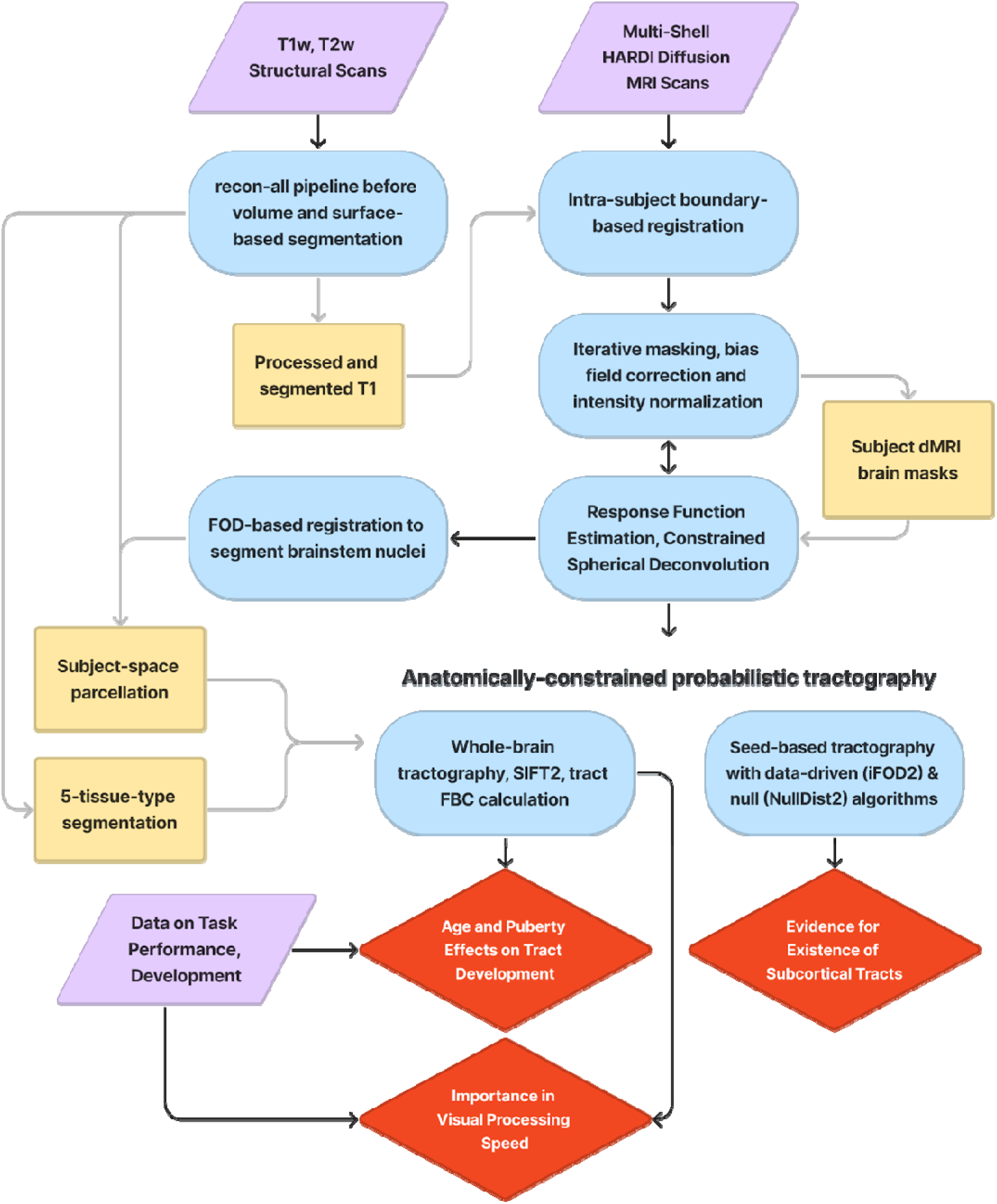
Analysis pipeline, including inputs (violet), processes (blue) intermediaries (yellow) and outputs pertinent to the three primary hypotheses (red). (FOD: Fibre Orientation Distribution; FBC: fibre bundle capacity, dMRI: diffusion MRI, T1w: T1-weighted images, T2w: T2-weighted images, SIFT: Spherical-deconvolution Informed Filtering of Tractograms)

Tract reconstruction was constrained based on participant neuroanatomy with anatomically constrained tractography (ACT), wherein streamlines were only allowed to traverse white and subcortical gray matter, and to terminate only in cortical or subcortical gray matter or at the base of the brain stem (Smith et al. 2012). ACT was guided by a tissue segmentation image generated using a combination of FreeSurfer cortical surface delineation, subcortical nuclei segmentation using FSL FIRST (Patenaude et al. 2011), the segmentation of hippocampal subfields, amygdalae, and brainstem with Freesurfer modules (Iglesias et al. 2015; Saygin et al. 2017) and other image processing operations (Smith et al. 2020). Brain stem nuclei segmented using the Brainstem Navigator probabilistic atlas (Bianciardi 2021) were subsequently introduced into these images in such a way that streamlines could either terminate within or pass through these nuclei, reflecting the uncertainty in which structural connections may synapse within them. Truncation of streamlines within subcortical nuclei was performed probabilistically in proportion to local fibre density to better reflect uncertainty in axonal penetration (Smith 2023). Streamline endpoints were assigned to gray matter ROIs using the radial search algorithm (Yeh et al. 2019); for brain stem nuclei, all streamlines intersecting each ROI were captured.

The set of cortical and subcortical tracts involved in visual information transfer were specified between ROIs selected based on previous research from our group (McFadyen, Mattingley, and Garrido 2019; McFadyen, Dolan, and Garrido 2020). All of these pathways are illustrated in **Figure 1**. The subcortical pathway consisted of connections between the superior colliculus and pulvinar (SC⍰Pul), superior colliculus and lateral geniculate nucleus (SC⍰LGN), superior colliculus and amygdala (SC⍰Amyg), and pulvinar and amygdala (Pul⍰Amyg); while the cortical pathway consisted of tracts connecting the lateral geniculate nucleus and V1 (LGN⍰V1), the primary visual area V1 and extrastriate cortices V2-V4 (V1⍰ES), extrastriate cortices and the inferotemporal cortex (ES⍰IT), and finally the amygdala and the inferotemporal cortex (Amyg⍰IT).

#### Targeted and null tractography

Objective assessment of evidence for the existence of each tract within the hypothesized subcortical pathway was performed by comparing the outcomes of targeted tractography against the corresponding null distribution, where tractography is not informed by the dMRI data. For simplicity, this test was performed exclusively using data from the first timepoint. For seeding from each of the regions involved in the subcortical visual amygdalar tracts (the amygdalae, pulvinar nuclei, and superior colliculi), 100,000 streamlines were generated using each of two tractography algorithms: iFOD2, and a corresponding “null distribution” algorithm where no fibre orientation information is used but streamlines nevertheless obey the same geometric integration constraints (Morris et al. 2008) and tissue segmentation. The null distribution tracking algorithm corresponding to iFOD2 is available in MRtrix3 as “tckgen -algorithm nulldist2”. Note that each tract could theoretically have a maximum of 200,000 streamlines assigned, since 100,000 streamlines were generated seeding from each of the two ROIs defining each tract.

#### Whole-brain tractography

To create whole-brain tractograms, ten million streamlines were generated for each participant’s dMRI image. “Dynamic seeding” was used for maximal compatibility with subsequent quantitative streamline tractography (Smith et al. 2015). Following whole-brain tractogram reconstruction, quantitative weights were ascribed to each streamline using the SIFT2 algorithm (Smith et al. 2015). For each tract of interest, the “fibre bundle capacity (FBC)” measure was quantified as the sum of weights of those streamlines ascribed to the pathway, scaled by the subject-specific proportionality coefficient within the SIFT model to ensure robust comparison across individuals (Smith et al. 2022).

### Developmental and Demographic Measures

Basic participant information included their sex (categorical; male and female), ethnicity (categorical; white, black, Hispanic, Asian and other), age at both timepoints, and the MRI scanner model (categorical; Siemens, General Electric, Phillips). Average scores on the Pubertal Development Scale (PDS) (Petersen, Crockett, Richards, & Boxer, 1988) were used to test associations between the structural connectivity of cortical and subcortical tracts and pubertal development. Specifically, we used scores from the parental version of the questionnaire as previously done (Thijssen, Collins & Luciana, 2020). The PDS measures pubertal development using three questions common to boys and girls, and two questions specific to each sex. Parental responses to the five questions were averaged, with possible scores ranging between 1 (pre-puberty) to 4 (post-puberty) (Petersen, Crockett, Richards, & Boxer, 1988).

### Processing Speed Measure

A measure of the speed of processing emotionally-salient human facial expressions was derived from response times in the 0-back condition (where working memory engagement is minimal) of the emotional visual N-Back task that participants performed during functional MRI scanning. Reaction times (RT) to images of faces with negative emotional expressions and faces with positive emotional expressions were averaged, and reaction time to images of places were used to control for overall individual differences in visual processing speed (Casey et al. 2018). We used response time data from the emotional N-Back task at both baseline and follow-up timepoints.

### Statistical Analyses

Statistical analyses were carried out using Python (version 3.9.12) and R software (version 4.3.0). FBC values from quantitative streamlines tractography were harmonized across scanners using the extensively validated Longitudinal ComBat method in R (Beer et al. 2020), with age, sex, and ethnicity included as covariates and subject and site included as random effects. FBC values were log-normalized prior to harmonization, since they were heavily right-skewed and log-normalization is known to improve ComBat harmonization for such data (Orlhac et al. 2022). Streamline counts from targeted tractography were not log-transformed or harmonized with ComBat, since these were positive count data and available implementations of ComBat cannot restrict the output to be positive integers.

The following models and tests were used to investigate the questions of interest:

1) The difference in streamline count between data-driven and null tractography algorithms as evidence for the existence of tracts.

Since streamline counts were overdispersed, negative binomial general linear models (GLMs) with the log link function were used. The main effect of interest was that of “Algorithm”, a binary variable with “Null” in the intercept such that the effect of using “iFOD2” (the other category) versus the null algorithm could be estimated. A model was fit for each tract independently, with fixed effects to control for the effects of scanner (since data were not harmonized between scanners), sex, and ethnicity. Only data from the first timepoint were used, and thus the effect of timepoint was not included in these models:

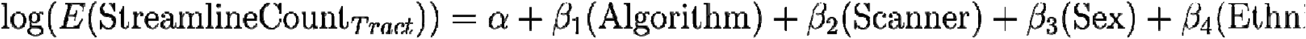

Linear models were used to test the other major hypotheses. These models all used log-normalized FBC values (unlike the previous GLMs which used streamline counts). All linear mixed-effects models controlled for the effects of sex and ethnicity (by including them as fixed effects). These models were fit only for tracts with evidence of existence based on the aforementioned analysis.

2) The importance of the structural connectivity of subcortical tracts for rapid processing of salient visual information, in this case images of human faces expressing positive or negative emotions

This model was fit once for each tract individually, and was used to test associations between tract structural connectivity and RT, and whether this association was different for emotional face vs. place stimuli. These models included each participant’s RT as the dependent variable, main effects for the structural connectivity of the tract (i.e., log-normalized FBC values) and stimulus category (emotional faces or places) as main effects, and the interaction term between structural connectivity and stimulus category to assess hypothesized stronger association between tract structural connectivity and RT to emotional faces than to places. The effect of timepoint (baseline vs follow-up) was also included in these models, since data were taken from both timepoints. Two-way and three-way interaction terms were also modelled between timepoint, structural connectivity and stimulus type, to capture potential differences in associations between structural connectivity and RT as a function of timepoint and stimulus type. Given each participant contributed multiple measurements for each model, as stimulus category and timepoint were within-subject factors, a random effect of participant was included in the intercept:

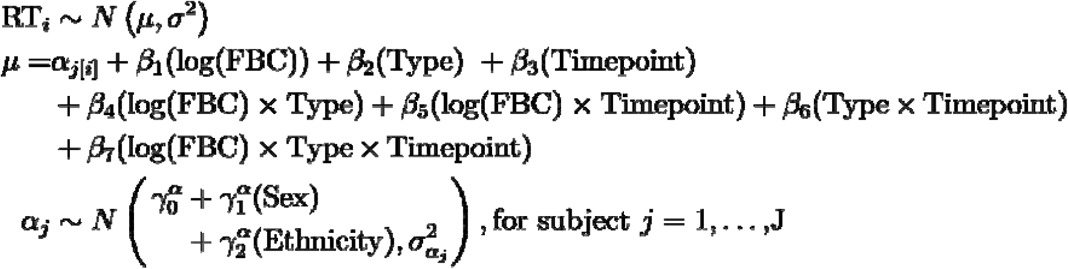

3) The developmental trajectory of visual tracts and the impact of puberty on their development, i.e., the longitudinal effects of age and puberty on their structural connectivity.

Change in the structural connectivity of each tract between the two timepoints was analyzed by including tract structural connectivity (i.e., log-transformed FBC of one subcortical or cortical tract) at the second timepoint as the dependent variable and structural connectivity at the first timepoint as a predictor. Given the collinearity of age and puberty (explored and visualized in **Supplementary Figure 1**) may complicate the estimation of both age and puberty effects in the same model, as well as potential sex differences in the effect of puberty, we compared three sets of models to select the final form of the model: a) models with only age effects; b) models with age and puberty effects; and c) models with age, puberty, and puberty-by-sex interaction effects (for details of the hierarchical model comparison process see **Supplementary Table 1**). The selected model (model b) modeled age and puberty at baseline, and the change in age and puberty between the two timepoints as fixed effects of interest:

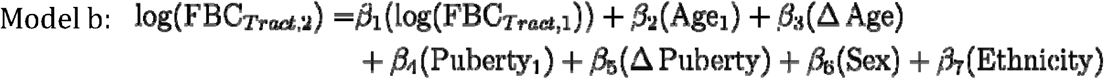

We first identified tracts whose structural connectivity was influenced by either age or puberty variables, using an F-test to compare the variance explained by model b and a null model that included no age or puberty variables) for each tract:

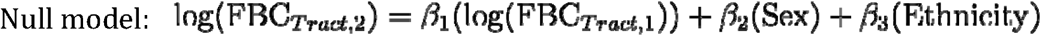

The significance of individual age- and puberty-related fixed effects was then tested for tracts where model comparison identified a significant difference to the null model.

4) The hypothesis that maturation trajectories may differ between the whole cortical and subcortical pathways.

Given that the models above were fit to each tract individually, they do not adequately address this more global hypothesis. To address such in a more tailored fashion, we constructed an exploratory model to investigate possible differences in the effects of age and puberty on structural connectivity of the cortical vs. subcortical pathways. This was done by collating all (log-transformed) FBC measures across tracts and defining a new binary “Pathway” factor indicating whether the FBC measurement belongs to a tract in the cortical or subcortical pathway. That model was defined with the interaction of this new binary variable with age and puberty fixed effects, to test whether these variables have a different impact on the change in structural connectivity of tracts in the subcortical pathway compared to those in the cortical pathway between baseline and follow-up timepoints. Differences in the mean structural connectivity of each tract were accounted for by including the categorical variable “Tract” in the model.

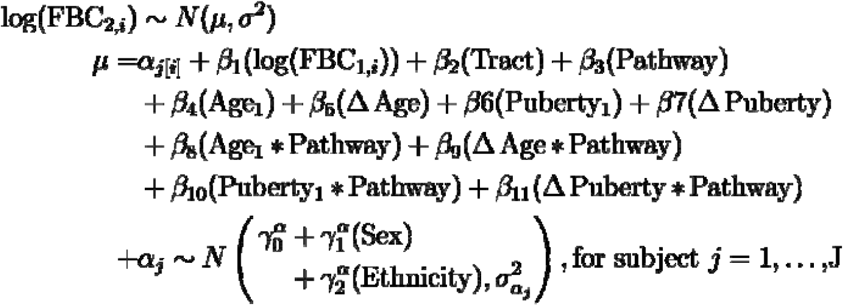

Since pathway interaction effects were significant (see the Results section), post-hoc versions of this model were used to determine the overall effect of puberty and age variables on structural connectivity separately for subcortical tracts and cortical tracts. Sex and ethnicity were entered as participant-level variables. Random effect of participant was included since, after collapsing FBC values across tracts, each participant contributes multiple measurements to the modeled data (as many measurements as there are tracts), with j and i respectively indexing participant and tract.

To control the family-wise error rate for any set of multiple analyses, p-values were corrected for false discovery rate (FDR) following the Benjamini-Hochberg procedure (Benjamini and Hochberg 1995).

## Results

We reconstructed every major tract in the cortical and subcortical visual pathways to the amygdala in the vast majority of participants across both time points using a whole-brain tractography algorithm. All tracts were reconstructed in more than 99% of participants, except for tracts between the right superior colliculus and right amygdala (93.3% of scans across both timepoints), left superior colliculus and left amygdala (97.2%), and right superior colliculus and right LGN (96.9%). Average streamline trajectories for tracts in the two pathways are presented visually in **Figure 3**; each depicted streamline represents one tract of one participant at one time point, where for each reconstructed tract for a participant, a mean spatial trajectory across the reconstructed streamlines was computed and transformed to template space.

**Figure 3:**
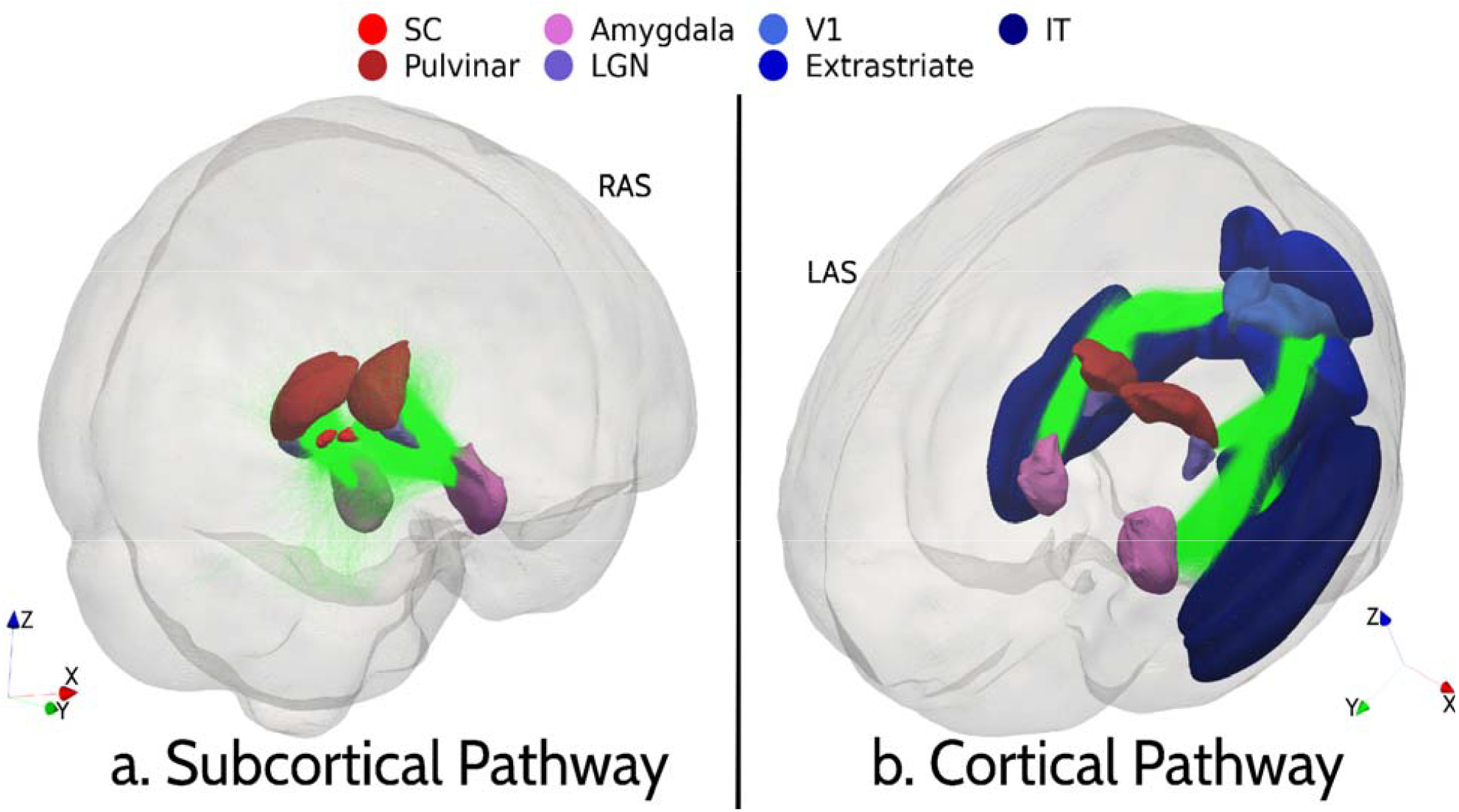
Regions of interest and tracts comprising the cortical and subcortical pathways, with each pathway shown in both hemispheres (subcortical pathway in panel a, cortical pathway in panel b). Each streamline shown represents the average trajectory for streamlines in one tract from one scan. Each region of interest is created by averaging regions of interest across participants. Tracts of interest in the subcortical pathway include the SC⍰Pul, SC⍰LGN, SC⍰Amyg and Pul⍰Amyg tracts, while tracts of interest in the cortical pathway include the V1⍰ES, LGN⍰V1, ES⍰IT and Amyg⍰IT tracts. Orientation markers: RAS, right anterior superior; LAS: left anterior superior. SC: superior colliculus; LGN: lateral geniculate nucleus; V1: primary visual cortex; Amyg: amygdala; IT: inferotemporal cortex; ES: extrastriate cortex.

### Evidence for the Existence of Subcortical Tracts

Negative binomial models were used to determine whether diffusion MRI data support the existence of subcortical tracts when compared against null distributions obtained with the null tractography algorithm. The list of the beta coefficients of the algorithm term in each GLM, indicating the difference of the data-driven and null tractography algorithms, is presented in **Table 2** (see **Supplementary Table 2** for complete model fitting results). All *p*-values in this and the following tables were FDR-corrected.

**Table 2:** Average streamline counts for each subcortical tract in each hemisphere, compared between the data-driven (iFOD2) and null tractography algorithms using negative binomial general linear model models.

| Tract | Null<br>*M* (*SD*) | iFOD2<br>*M* (*SD*) | Effect<br>size | *SE* | *z*<br>value | *p*<sub>FDR</sub> |
| --- | --- | --- | --- | --- | --- | --- |
| r.SC↔Pul | 22169 (5332) | 4811 (5132) | -1.56 | 0.016 | -97.4 | 0 |
| l.SC↔Pul | 23339 (5758) | 4900 (5208) | -1.58 | 0.015 | -102.5 | 0 |
| **r.SC↔LGN** | **3234 (824)** | **4252 (4943)** | **0.26** | **0.019** | **13.5** | **9.3e-41** |
| **l.SC↔LGN** | **3201 (800)** | **4126 (4252)** | **0.25** | **0.017** | **14.8** | **9.6e-49** |
| r.SC↔Amyg | 1015.2 (376) | 186 (308) | -1.72 | 0.019 | -88.5 | 0 |
| l.SC↔Amyg | 1152 (395) | 319 (504) | -1.30 | 0.019 | -70.3 | 0 |
| **r.Pul↔Amyg** | **372 (119)** | **6255 (3897)** | **2.81** | **0.010** | **285.3** | **0** |
| **l.Pul↔Amyg** | **382 (120)** | **6432 (4234)** | **2.82** | **0.010** | **275.9** | **0** |

*Note.* *p*-values that are too small to accurately report due to software constraints (smaller than $\sim 1e-300$ ) are reported as 0. Rows where data-driven tractography leads to significantly more streamlines than null tractography are highlighted in bold. All rows had significant differences between the two algorithms. SC: superior colliculus; LGN: lateral geniculate nucleus; Pul: pulvinar; Amyg: amygdala; r: right; l: left; FDR: false discovery rate.

**Table 2:**
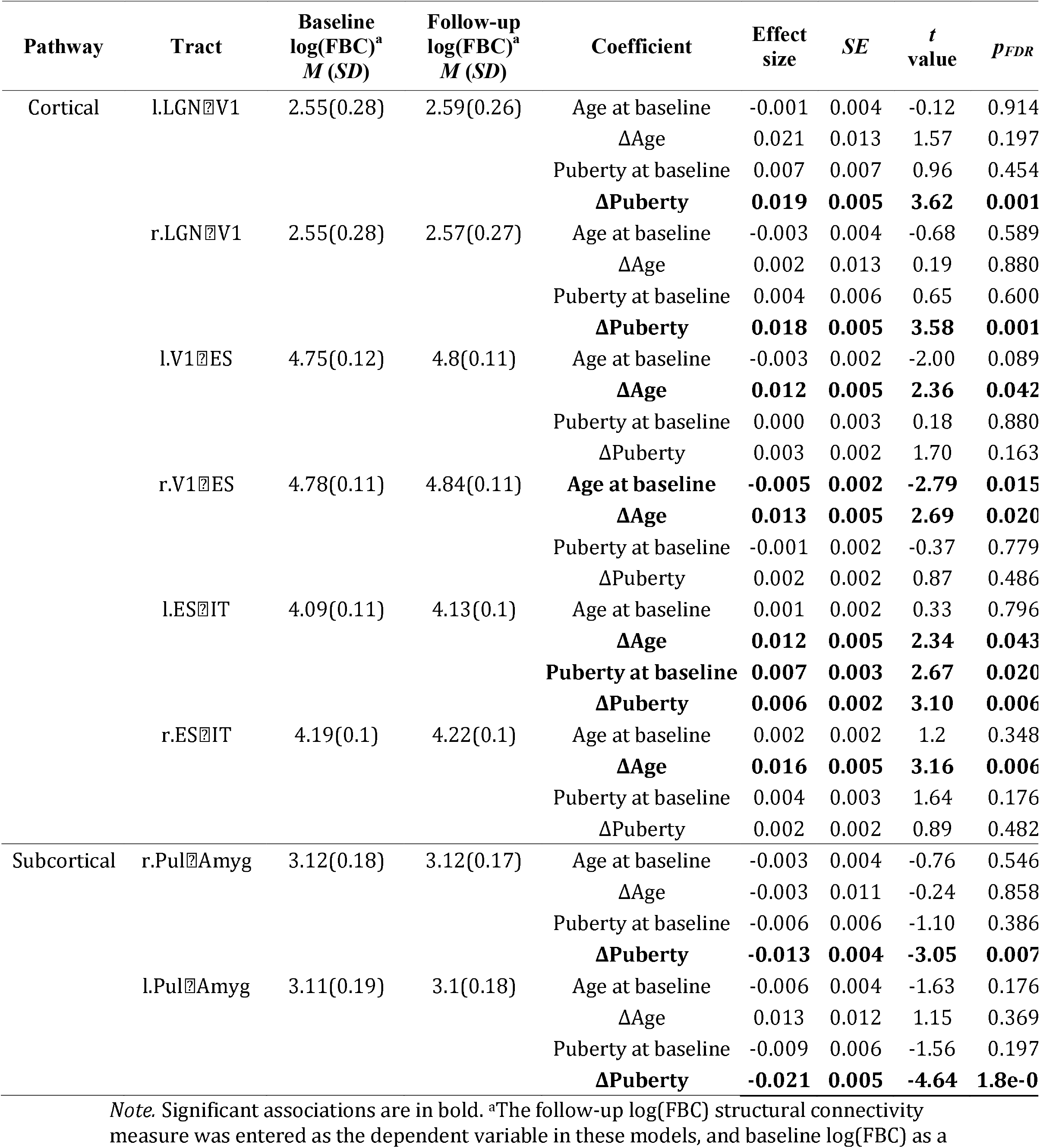
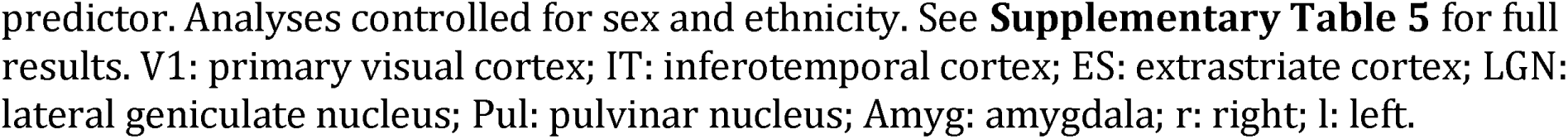
Association between change in tract structural connectivity, defined as the logtransformed Fibre Bundle Capacity (FBC), between baseline and follow-up and baseline age, change in age (follow-up minus baseline), baseline pubertal development, and change in pubertal development.

| Pathway | Tract | Baseline<br>log(FBC)<sup>a</sup><br>M (SD) | Follow-up<br>log(FBC)<sup>a</sup><br>M (SD) | Coefficient | Effect<br>size | SE | t<br>value | p<sub>FDR</sub> |
| --- | --- | --- | --- | --- | --- | --- | --- | --- |

| Pathway | Tract | Baseline<br>log(FBC)<sup>a</sup><br>*M (SD)* | Follow-up<br>log(FBC)<sup>a</sup><br>*M (SD)* | Coefficient | Effect<br>size | *SE* | *t*<br>value | *p*<sub>FDR</sub> |
| --- | --- | --- | --- | --- | --- | --- | --- | --- |
| Cortical | l.LGN↔V1 | 2.55(0.28) | 2.59(0.26) | Age at baseline | -0.001 | 0.004 | -0.12 | 0.914 |
|  |  |  |  | ΔAge | 0.021 | 0.013 | 1.57 | 0.197 |
|  |  |  |  | Puberty at baseline | 0.007 | 0.007 | 0.96 | 0.454 |
|  |  |  |  | **ΔPuberty** | **0.019** | **0.005** | **3.62** | **0.001** |
|  | r.LGN↔V1 | 2.55(0.28) | 2.57(0.27) | Age at baseline | -0.003 | 0.004 | -0.68 | 0.589 |
|  |  |  |  | ΔAge | 0.002 | 0.013 | 0.19 | 0.880 |
|  |  |  |  | Puberty at baseline | 0.004 | 0.006 | 0.65 | 0.600 |
|  |  |  |  | **ΔPuberty** | **0.018** | **0.005** | **3.58** | **0.001** |
|  | l.V1↔ES | 4.75(0.12) | 4.8(0.11) | Age at baseline | -0.003 | 0.002 | -2.00 | 0.089 |
|  |  |  |  | **ΔAge** | **0.012** | **0.005** | **2.36** | **0.042** |
|  |  |  |  | Puberty at baseline | 0.000 | 0.003 | 0.18 | 0.880 |
|  |  |  |  | ΔPuberty | 0.003 | 0.002 | 1.70 | 0.163 |
|  | r.V1↔ES | 4.78(0.11) | 4.84(0.11) | **Age at baseline** | **-0.005** | **0.002** | **-2.79** | **0.015** |
|  |  |  |  | **ΔAge** | **0.013** | **0.005** | **2.69** | **0.020** |
|  |  |  |  | Puberty at baseline | -0.001 | 0.002 | -0.37 | 0.779 |
|  |  |  |  | ΔPuberty | 0.002 | 0.002 | 0.87 | 0.486 |
|  | l.ES↔IT | 4.09(0.11) | 4.13(0.1) | Age at baseline | 0.001 | 0.002 | 0.33 | 0.796 |
|  |  |  |  | **ΔAge** | **0.012** | **0.005** | **2.34** | **0.043** |
|  |  |  |  | **Puberty at baseline** | **0.007** | **0.003** | **2.67** | **0.020** |
|  |  |  |  | **ΔPuberty** | **0.006** | **0.002** | **3.10** | **0.006** |
|  | r.ES↔IT | 4.19(0.1) | 4.22(0.1) | Age at baseline | 0.002 | 0.002 | 1.2 | 0.348 |
|  |  |  |  | **ΔAge** | **0.016** | **0.005** | **3.16** | **0.006** |
|  |  |  |  | Puberty at baseline | 0.004 | 0.003 | 1.64 | 0.176 |
|  |  |  |  | ΔPuberty | 0.002 | 0.002 | 0.89 | 0.482 |
| Subcortical | r.Pul↔Amyg | 3.12(0.18) | 3.12(0.17) | Age at baseline | -0.003 | 0.004 | -0.76 | 0.546 |
|  |  |  |  | ΔAge | -0.003 | 0.011 | -0.24 | 0.858 |
|  |  |  |  | Puberty at baseline | -0.006 | 0.006 | -1.10 | 0.386 |
|  |  |  |  | **ΔPuberty** | **-0.013** | **0.004** | **-3.05** | **0.007** |
|  | l.Pul↔Amyg | 3.11(0.19) | 3.1(0.18) | Age at baseline | -0.006 | 0.004 | -1.63 | 0.176 |
|  |  |  |  | ΔAge | 0.013 | 0.012 | 1.15 | 0.369 |
|  |  |  |  | Puberty at baseline | -0.009 | 0.006 | -1.56 | 0.197 |
|  |  |  |  | **ΔPuberty** | **-0.021** | **0.005** | **-4.64** | **1.8e-0** |

*Note.* Significant associations are in bold. <sup>a</sup>The follow-up log(FBC) structural connectivity measure was entered as the dependent variable in these models, and baseline log(FBC) as a

Considering the estimated effect sizes, the natural logarithm of the number of streamlines generated based on Fibre Orientation Distributions (FOD) derived from the diffusion signal is 0.26 and 0.25 greater than would be expected by chance for the right and left superior colliculus-pulvinar tracts respectively, and 2.81 and 2.82 greater for both the right and the left pulvinar-amygdala tracts respectively. Streamline counts for each subcortical tract resulting from either algorithm (with seeding from both ends of each tract) are presented in **Figure 4**. For some tracts, anatomically-constrained null tractography produced more streamlines (bilateral SC⍰amygdala and bilateral SC⍰pulvinar).

**Figure 4:**
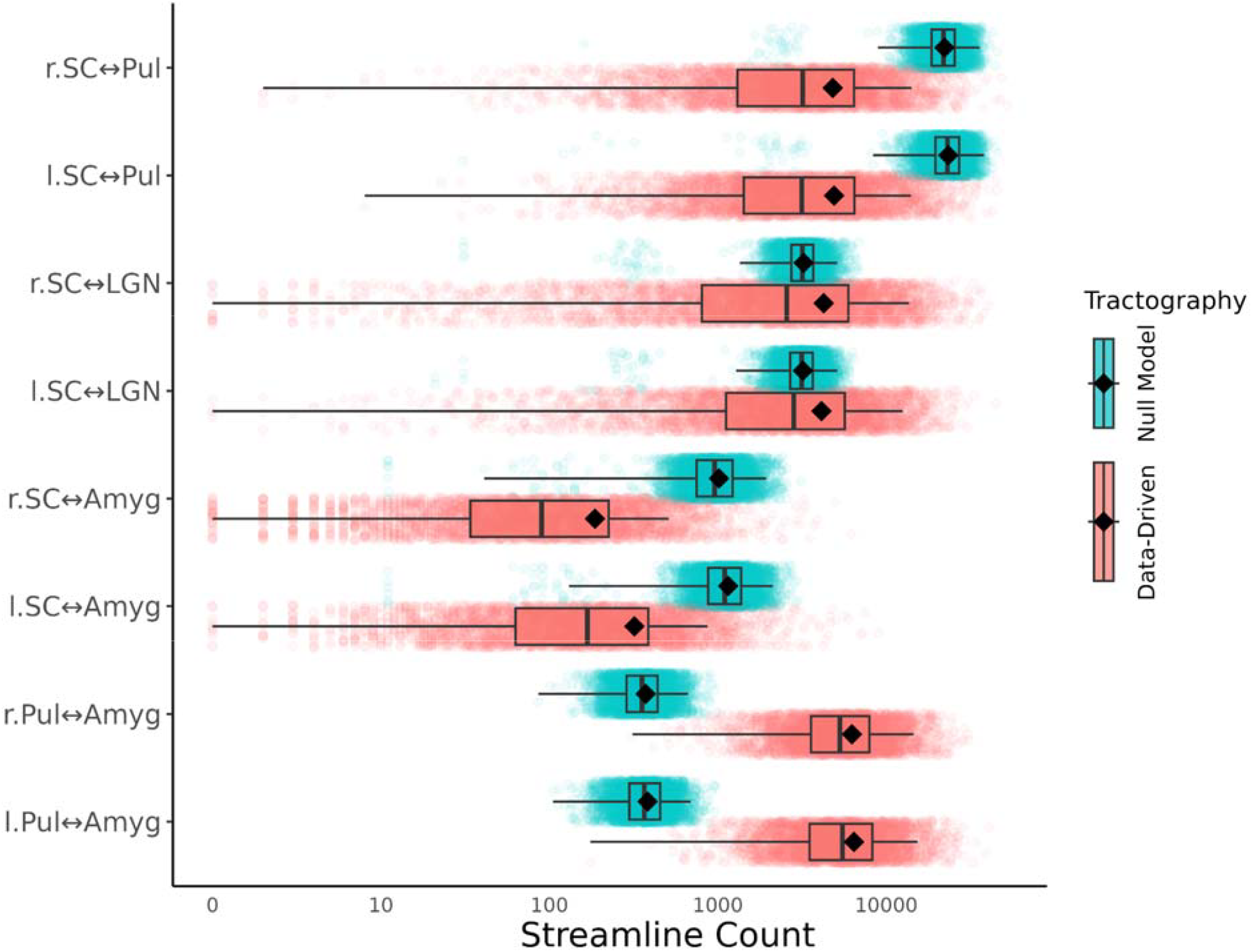
Tract streamline counts for subcortical tracts comparing FOD-based tracking to an equivalent null tractography algorithm. For each algorithm, 100,000 streamlines were seeded from each region; the streamline count for each tract is the sum of streamlines seeded from either region that intersected the other. Boxplots represent the first, second and third quartiles, with whiskers extending for 1.5 * interquartile range from the first and third quartile. Black diamonds represent the mean streamline count SC: superior colliculus; LGN: lateral geniculate nucleus; Pul: pulvinar nucleus; Amyg: amygdala; r: right; l: left.

Based on these results, the left and right pulvinar⍰amygdala connections were included in further analyses as representing the subcortical pathway. Left and right SC-LGN connections were not analysed further since the evidence for their existence was much weaker, previous literature on their importance in the subcortical pathway is mixed, and neither of their two hypothesized downstream tracts (SC⍰pulvinar and SC⍰amygdala) had evidence of existence in these data.

### Greater Subcortical and Cortical Structural Connectivity are Associated with Faster Processing Speed for Faces and Places

It has been hypothesized that the subcortical pathway facilitates rapid processing of emotional visual stimuli. Here, we used data from both timepoints to test whether subcortical or cortical structural connectivity were associated with the speed of responding in the 0-Back trials of the emotional visual N-back task, and whether this association was specific to emotional face stimuli, when compared to place stimuli.

The structural connectivity (log-transformed FBC) of the right pulvinar⍰amygdala tract was negatively associated with RT in 0-Back trials of the emotional visual N-Back task (⍰ =-33.08, *p*_*FDR*_ = 0.006); that is, participants with stronger right pulvinar-amygdala connections responded relatively faster than those with weaker connectivity in that tract. However, the interaction of this effect with stimulus category (emotional faces vs. places) was not significant. The structural connectivity of three cortical tracts had a similarly negative and non-specific associations with response time: right V1⍰ES (⍰ = -52.58, *p*_*FDR*_ = 0.005), left V1⍰ES (⍰ = -77.00, *p*_*FDR*_ < 0.001), and left ES⍰IT (⍰ = -48.03, *p*_*FDR*_ = 0.021). Indeed, neither the connectivity-by-stimulus category interaction effects nor the three-way interaction effects (connectivity-by-timepoint-by-category) were significant for any tract. Overall, this indicates that whilst greater connectivity was associated with faster processing speed, this was a general effect observed regardless of whether stimuli were emotional faces or places. An additional result was that the interaction term between structural connectivity and timepoint was significant and positive only for pulvinar⍰amygdala tracts in both the right (⍰ = 56.66, *p*_*FDR*_ < 0.001) and left (⍰ = 33.77, *p*_*FDR*_ = 0.020) hemispheres, indicating that the relationship between connectivity and response time was more negative at baseline than at the second timepoint (see **Supplementary Table 3** for model fitting results).

### Developmental Changes in Structural Connectivity

Next, we investigated changes in structural connectivity across baseline and follow-up. Model comparison revealed that for most subcortical and cortical tracts, linear models that included main effects of age and puberty explained significantly more variance than models without these effects, suggesting developmental changes in structural connectivity of these tracts during the studied age window (**Supplementary Table 4)**. Testing the significance of individual effects (for tracts where developmental model comparison with the null was significant) revealed that a longer time between timepoints (ΔAge) was positively associated with FBC at timepoint 2 in bilateral V1⍰ES tracts and ES⍰IT tracts (**Table 2, Figure 5**). Hence, participants whose chronological age increased more between timepoints, when controlling for pubertal development, showed increased bilateral V1⍰ES and ES⍰IT structural connectivity (see **Supplementary Table 5** for complete model fitting results).

**Figure 5:**
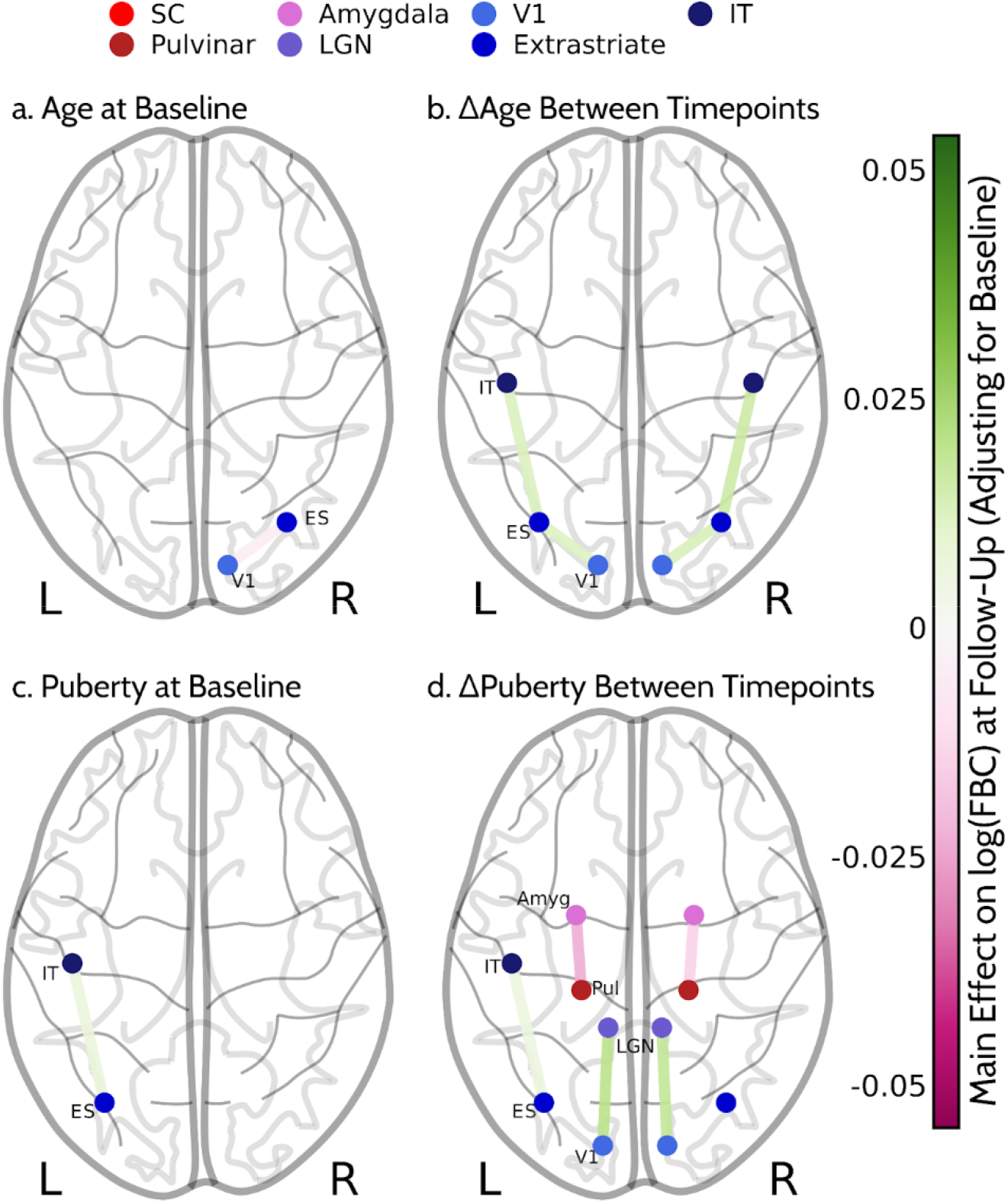
Association between change in tract structural connectivity, defined as the log-transformed Fibre Bundle Capacity (FBC), between baseline and follow-up and a) baseline age, b) change in age (follow-up minus baseline), c) baseline pubertal development, and d) change in pubertal development. Only significant associations are shown. SC: superior colliculus; Pul: pulvinar nucleus; LGN: lateral geniculate nucleus; Amyg: amygdala; ES: extrastriate cortex; IT: inferotemporal cortex, R: right, L: left.

Change in parent-reported pubertal development was similarly positively associated with FBC in bilateral LGN⍰V1 tracts and left ES⍰IT tract but was negatively associated with FBC in subcortical bilateral pulvinar⍰amygdala tracts (see **Table 2, Figure 6**). This suggests that participants with greater pubertal progression between timepoints showed increased bilateral LGN⍰V1 and left ES⍰IT structural connectivity, but decreased Pul⍰Amyg connectivity, after controlling for age.

The main effects of baseline age and puberty were sparser. Pubertal development at baseline was positively associated with left ES⍰IT connectivity while age at baseline was negatively associated with right ES⍰V1 connectivity. Overall, this suggests that pubertal development is associated with weakening of the subcortical pathway while chronological age and pubertal development are associated with strengthening of the cortical visual pathway.

### Differential trajectories of cortical and subcortical pathways

Next, we ran an exploratory linear mixed-effects model of the interaction between age/puberty and tract pathway membership (i.e., whether a structural connectivity measurement is from a tract in the subcortical pathway or one in the cortical pathway). We found that membership of the subcortical pathway significantly interacted with ΔPuberty (*p*_*FDR*_ = 2.5e-21), baseline puberty (*p*_*FDR*_ = 0.002) and baseline age (*p*_*FDR*_ = 0.031). Post-hoc models of the separate effects of age and puberty variables on structural connectivity in either pathway revealed significant positive association with ΔAge (*β* = 0.013, *p*_*FDR*_ = 0.005) and ΔPuberty (⍰ = 0.008, *p*_*FDR*_ = 3.1e-6) for the cortical pathway and a significant negative association with ΔPuberty for the subcortical pathway (⍰ = -0.018, *p*_*FDR*_ = 4.2e-6) (**Table 3**, see **Supplementary Tables 6** for detailed model fitting results).

**Table 3:** Result of the linear mixed model analysis of change in the structural connectivity in cortical and subcortical pathways between baseline and follow-up timepoints as a function of pubertal development and chronological age, combined across tracts.

| Pathway | Coefficient | Estimate | SE | df | t value | $p_{FDR}$ |
| --- | --- | --- | --- | --- | --- | --- |
| Cortical | Age at baseline | -0.001 | 0.001 | 4287.9 | -1.08 | 0.459 |
| | ΔAge | **0.013** | **0.004** | **4275.5** | **2.98** | **0.005** |
|  | Puberty at baseline | 0.004 | 0.002 | 4275.9 | 1.67 | 0.121 |
| | ΔPuberty | **0.008** | **0.002** | **4275.4** | **4.88** | **3.1e-6** |
| Subcortical | Age at baseline | -0.004 | 0.003 | 4248.5 | -1.36 | 0.195 |
| | ΔAge | 0.005 | 0.009 | 4245.2 | 0.49 | 0.625 |
|  | Puberty at baseline | -0.008 | 0.005 | 4243.8 | -1.70 | 0.121 |
| | ΔPuberty | **-0.018** | **0.004** | **4243.6** | **-4.80** | **4.2e-6** |

*Note*. Analyses controlled for sex and ethnicity, and random intercepts were included for participants.

## Discussion

The present study is the largest investigation of visual subcortical amygdalar tracts to date and the first assessment of their developmental trajectory in early adolescence. We found corroborative evidence for the existence of major bilateral tracts in the subcortical pathway to the amygdala in young adolescents, and demonstrate that pulvinar-amygdala connectivity strength, a major segment of the so-called “low-road”, is linked to faster processing of visual stimuli. However, this association was not specific to emotional face stimuli, contrary to what was expected; nor was it specific to the subcortical route, as response speed was similarly associated with stronger cortical connectivity. Further, we show divergent maturation patterns for the subcortical and cortical pathways over development: while subcortical connectivity to the amygdala decreased during pubertal development, connectivity in the visual cortical pathway increased with pubertal development and chronological age.

### Evidence for the Existence of a Subcortical Pathway to Amygdala

As the lack of direct anatomical evidence has led to controversies about the existence and function of a subcortical pathway to amygdala in humans (Pessoa and Adolphs 2010), we first compared our streamline reconstruction of each tract of interest against a corresponding null model where fibre orientation information from dMRI was not used (Morris et al., 2008). Tract reconstruction studies do not usually contrast their result with appropriate null models, considered requisite for discovery reporting in any other context. Whilst we did not find evidence that the superior colliculus-pulvinar or superior colliculus-amygdala pathways could be reconstructed above what would be expected based on anatomic geometry alone (in contrast to Rafal et al., 2015; Tamietto et al., 2012 but they did not use null tractography), we found very convincing evidence for the bilateral pulvinar-amygdala tracts and bilateral tracts linking the superior colliculi and lateral geniculate nuclei. These findings are concordant with evidence from optogenetics, single-neuron electrophysiology, and axon tracing from rodents and non-human primates (Elorette et al. 2018; Inagaki et al. 2022; Linke et al. 1999). Furthermore, our results align with investigations of the subcortical visual pathway in humans using functional imaging with positron emission tomography (Morris, Öhman, and Dolan 1999), functional MRI (Kragel et al. 2021; McFadyen, Mattingley, and Garrido 2019), and magnetoencephalography (Garrido et al. 2012; Garvert et al. 2014; McFadyen et al., 2017), as well as evidence from dMRI studies using alternative methodologies (Huang et al. 2022b; Rafal et al. 2015; Tamietto et al. 2012, McFadyen et al., 2019).

Our study differs from most earlier diffusion MRI tractography investigations in several important respects. First, many tractography studies of the subcortical pathway have used deterministic tractography, where high precision can give a misleading sense of accuracy. Here, we used probabilistic tractography not only as this is intrinsic to comparison against a corresponding tractography null model but also since it is generally better able to capture the underlying neuroanatomy, especially in subcortical regions (Maffei et al. 2022). Most previous studies performed streamline tracking based on the diffusion tensor model, which has known limitations in resolving intra-voxel fibre orientations and complex fibre geometries and may lead to the extensive reconstruction of false positives. To address these limitations, we used FODs derived with the more modern constrained spherical deconvolution approach (Jeurissen et al. 2014). This echoes a previous study of the human connectome project data that also found evidence for tracts between the superior colliculus, pulvinar and amygdala (McFadyen, Mattingley, and Garrido 2019). Here, we refined this methodological approach by incorporating the Anatomically-Constrained Tractography framework, ensuring that streamlines only traversed and terminated where anatomically plausible (Smith et al. 2012; Smith et al. 2020). Moreover, we mapped all regions of interest using validated probabilistic atlases (García-Gomar et al. 2019; Wang et al. 2015), in contrast to the manually drawn superior colliculus masks used in McFadyen et al. (2019). Additionally, we investigated these pathways in a young population (9–12 years of age) and the maturation of these pathways during early pubertal development using longitudinal data.

### Stronger Structural Connectivity Correlates with Faster Responses to Visual Stimuli

Besides evidence for the likely existence of pulvinar-amygdala tracts, we show that their structural connectivity strength is associated with faster responses on a visual task. Given the hypothesized role of the subcortical pathway in the rapid processing of emotionally salient stimuli (Kalhan et al., 2022; McFadyen et al., 2017; Vuilleumier et al., 2003), we expected such relationship between response speed and structural connectivity to be specific to emotionally face stimuli. However, the association between structural connectivity and RT was seen across both stimulus categories (emotional faces and places). It is worth noting differences between tasks and measures used in the literature, which may explain the seemingly inconsistent findings. Firstly, faster response times to fearful than neutral faces were previously associated with stronger amygdala activity (Kalhan et al., 2022), but that study did not investigate amygdala structural connectivity with the pulvinar. Hence, it is possible that such hastening was facilitated via a different tract. Secondly, the task completed by participants used continuous flash suppression, whereby faces slowly emerged into consciousness and participants were asked to press a button as soon as they saw a face. This is a type of detection task and a different process to the one studied in this paper, whereby participants were asked to match the current stimulus with a previously seen stimulus. In McFadyen et al., (2019) an association was found between structural connectivity of the pulvinar-amygdala tract and performance in a task requiring emotion recognition, but with performance quantified as *accuracy* rather than *response speed*. Again, that task engaged a different process to the stimulus matching task considered here, as overall stimuli characteristics needed to be attended to, rather than specifically the emotion pictured. Attentionally-demanding tasks, such as the one used in the ABCD Study might suppress amygdalar responses to emotional stimuli (Pessoa et al., 2002; Williams et al., 2005), possibly via suppression of the subcortical pathway, which might explain the lack of an emotional faces-specific effect in the association with RT and structural connectivity. Another possibility is that the subcortical pathway is more broadly involved in visual processing than often assumed. Indeed, there is evidence that the subcortical amygdala pathway carries information for non-emotional stimuli like neutral faces (McFadyen et al., 2017) and images of food (Sato et al., 2019).

While we had predicted that hastening of behavioral responses would be enabled by stronger structural connectivity specifically in the subcortical amygdala pathway, this association was also seen in some cortical tracts—namely bilateral V1-ES and left ES-IT— suggesting that this speed advantage is general to stronger connectivity in both subcortical and cortical tracts. In contrast, in McFadyen et al. (2017), data-driven neural simulations for magnetoencephalography revealed a clear temporal advantage of the subcortical connection over the cortical connection in influencing amygdala activity. However, that study used an incidental gender discrimination task and did not investigate the behavioural relevance of such temporal advantage. Nonetheless, it is still plausible that the speed of behavioral responses to salient stimuli (Kalhan et al., 2022; Koller et al., 2019) is facilitated by pulvinar-amygdala connectivity not just because of its stronger connectivity but also due to its shorter length relative to the cortical pathway to amygdala.

While similar associations between subcortical and cortical structural connectivity and response time to visual stimuli were observed across timepoints, only the subcortical pathway showed a greater association between stronger structural pulvinar-amygdala connectivity bilaterally and faster response speed at baseline (age 9-10 years old) than follow-up (age 11-12). This pattern of results suggests that the functional role of the subcortical tracts to amygdala may lessen over development, which aligns with the developmental results discussed below.

### Opposite Developmental Trajectories of the Cortical and Subcortical Pathways

We found diverging developmental trajectories for the cortical and subcortical tracts across normative development in early adolescence. Specifically, the subcortical pulvinar-amygdala pathway tracts weakened bilaterally during early adolescence with pubertal development, while some of the cortical tracts strengthened with pubertal development and chronological age. Exploratory analyses for overall pathway effects confirmed the existence of these two broad, opposite trajectories for the two pathways across puberty. These results are consistent with the finding in a different US longitudinal cohort showing that amygdala volume changes during adolescence correlate with both pubertal development and chronological age (Goddings et al., 2014).

To the best of our knowledge, this weakening trajectory has not previously been demonstrated in humans or animal models. This is partly because the existing literature on the visual subcortical pathway has focused largely on its reconstruction, and functional and behavioral roles, rather than its maturation across development. Indeed, the small number of developmental studies of visual pathways have investigated maturation of either the major (cortical) tracts or across the whole brain (i.e., globally rather than on the level of individual tracts) (Goddings et al. 2021; Lynch et al. 2020; Lebel, Treit, and Beaulieu 2019). While similar patterns have been noted in major amygdalar connections, such as the amygdalofugal pathway and the anterior commissure (Azad et al. 2021; Goetschius et al. 2019), the literature on the development of visual tracts to the amygdala remains scant.

One potential suggested role for the subcortical amygdala pathway has been in responding to visual stimuli from very early on in life, for example in early neonatal responses to impending collision (Ball and Tronick 1971). As with other visual pulvinar circuitry (notably pulvinar to the motion sensitive area, hMT+/V5), the subcortical pathway to the amygdala may provide a route for emotionally salient visual information that rapidly develops early in development, and then weakens as the canonical pathways for visual information mature (Warner et al. 2015; Bourne and Morrone 2017). This would be in keeping with the known selective pruning of connectivity in the developing human brain, particularly through childhood and adolescence (Goddings et al. 2019; Tymofiyeva et al. 2014). Interestingly, studies of blindsight suggest that the subcortical pathway may remain plastic and adaptively myelinate if the cortical pathway is damaged (Tamietto et al. 2012). This, together with our findings of divergent maturation patterns for subcortical and cortical amygdala pathways, suggests that subcortical and cortical pathways may weaken and strengthen, respectively.

An alternative explanation may be that the fine subcortical pathways prune simply as part of a broader tendency for the structural connectome to become less globally integrated and more modularized (Kaiser 2017; Tymofiyeva et al. 2014). As the connectome matures, weaker auxiliary connections that provide alternative communication routes between neural modules may be eliminated as a sparser connectome develops, and this *general* tendency (rather than a pruning mechanism *specific* to the subcortical pathway) may account for our observations.

### Limitations

The ABCD data used here came from participants aged between 9 and 12 years old. While the longitudinal aspect of this study is a strength when studying development, only two timepoints were available when the analyses were carried out. Future work could incorporate later timepoints, capturing structural connectivity changes in later pubertal development. Further, considering the proposed importance of the subcortical pathway in early development, future work could investigate developmental changes in structural connectivity of subcortical and cortical pathways in early childhood, a developmental period not captured in this study.

We used a parental survey as a proxy for pubertal development, consistently with previous research using the ABCD study data (Thijssen, Collins & Luciana, 2019). Future studies could consider more objective measures of physiological puberty, although these can be difficult to acquire at scale.

The ABCD study is a multisite study, which allowed a large sample size. All analyses controlled for data collection site, including explicit harmonization of FBC values across scanners. Such harmonization was however not possible for the streamline count data utilized in initial tests for the existence of connections.

## Conclusion

Here, we reconstructed the cortical and subcortical visual pathways to the amygdala in the largest sample to date. We add direct evidence for the existence a subcortical pathway and demonstrate association between its connectivity strength and faster response to visual stimuli regardless of its emotional content. While a similar association was seen with cortical connectivity, only the subcortical pathway showed a negative association between structural connectivity strength and response speed between timepoints. Further, using longitudinal data, we demonstrate that the cortical and subcortical amygdala pathways have distinct maturation trajectories across early adolescence, with cortical pathways strengthening and subcortical weakening with pubertal development.

