## Supplementary Materials for "Weakening of subcortical and strengthening of cortical visual pathways across early adolescence"

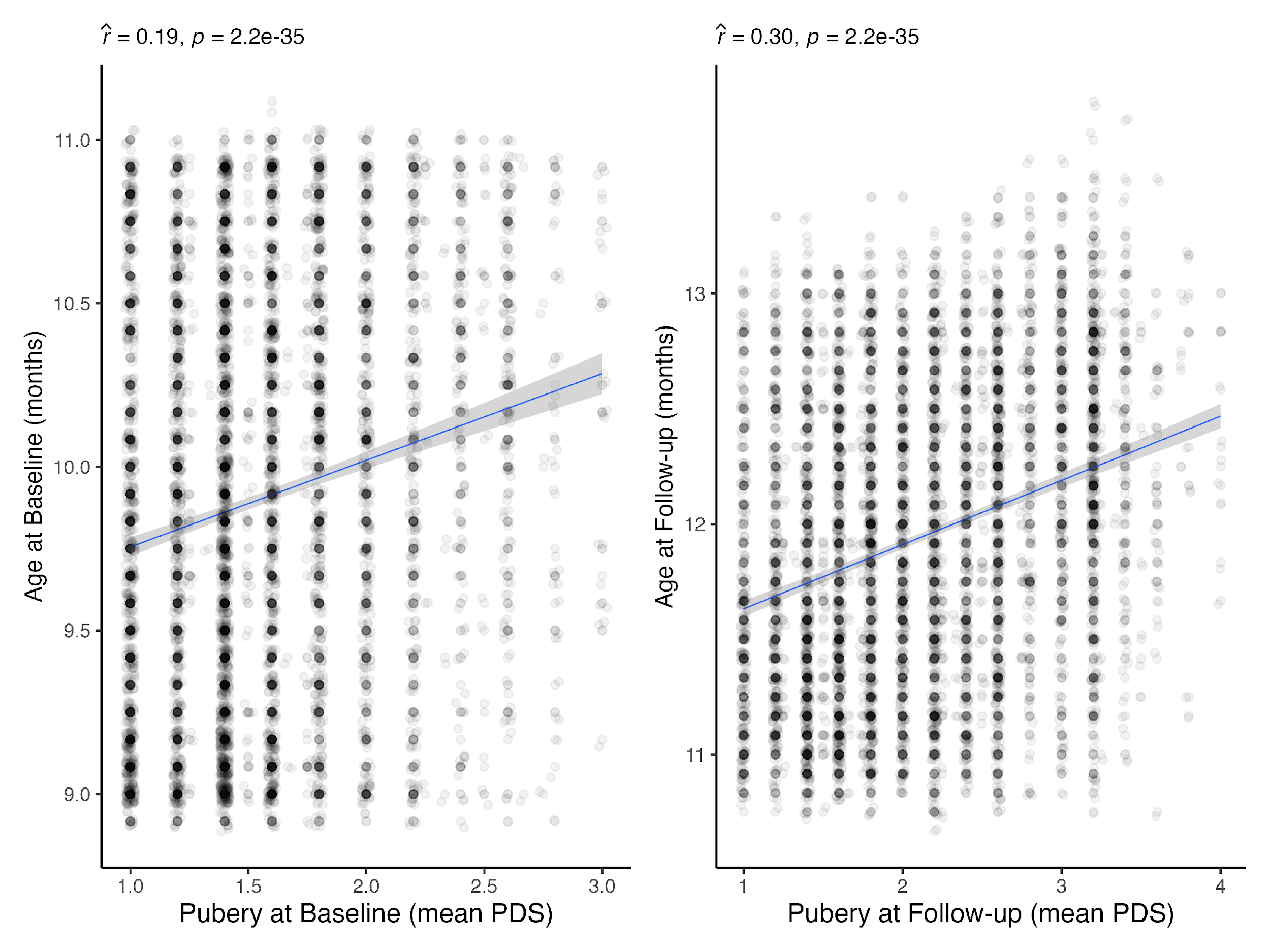

***Supplementary Figure 1:*** *Exploration of collinearity (using Pearson’s correlation tests) between age and puberty at the baseline and follow-up timepoints.*

***Supplementary Table 1:*** *Model selection results. For each tract, results of fitting each of the three models are presented for the variables of interest. The results of hierarchical F tests are presented for each tract as well. As can be seen, “Puberty” models explained significantly greater variance than the “Base” models across several tracts, while the “Interaction” models did not.*

| Pathway | Tract | Model | Coefficient | Effect size | SE | *t* value | *p* (Model Term) | *p* (Base vs. Puberty) | *p* (Puberty vs. Interaction) |
| --- | --- | --- | --- | --- | --- | --- | --- | --- | --- |
| Subcortical | r.Pul↔Amyg | Base | Age_1 | -0.007 | 0.003 | -1.93 | 0.053 | 0.009 | 0.394 |
|  |  | Base | ΔAge | -0.007 | 0.011 | -0.67 | 0.501 |  |  |
|  |  | Puberty | Age_1 | -0.003 | 0.004 | -0.76 | 0.447 |  |  |
|  |  | Puberty | ΔAge | -0.003 | 0.011 | -0.24 | 0.809 |  |  |
|  |  | Puberty | Puberty_1 | -0.006 | 0.006 | -1.10 | 0.272 |  |  |
|  |  | Puberty | ΔPuberty | -0.013 | 0.004 | -3.05 | 0.002 |  |  |
|  |  | Interaction | Age_1 | -0.003 | 0.004 | -0.72 | 0.469 |  |  |
|  |  | Interaction | ΔAge | -0.003 | 0.011 | -0.26 | 0.797 |  |  |
|  |  | Interaction | Puberty_1 | -0.006 | 0.006 | -1.12 | 0.264 |  |  |
|  |  | Interaction | ΔPuberty | -0.010 | 0.006 | -1.68 | 0.094 |  |  |
|  |  | Interaction | ΔPuberty:SexMale | -0.007 | 0.008 | -0.85 | 0.394 |  |  |
|  | l.Pul↔Amyg | Base | Age_1 | -0.012 | 0.004 | -3.42 | 6.2e-04 | 1.9e-05 | 0.136 |
|  |  | Base | ΔAge | 0.006 | 0.011 | 0.50 | 0.617 |  |  |
|  |  | Puberty | Age_1 | -0.006 | 0.004 | -1.63 | 0.102 |  |  |
|  |  | Puberty | ΔAge | 0.013 | 0.012 | 1.15 | 0.252 |  |  |
|  |  | Puberty | Puberty_1 | -0.009 | 0.006 | -1.56 | 0.119 |  |  |
|  |  | Puberty | ΔPuberty | -0.021 | 0.005 | -4.64 | 3.7e-06 |  |  |
|  |  | Interaction | Age_1 | -0.006 | 0.004 | -1.57 | 0.117 |  |  |
|  |  | Interaction | ΔAge | 0.013 | 0.012 | 1.12 | 0.264 |  |  |
|  |  | Interaction | Puberty_1 | -0.009 | 0.006 | -1.59 | 0.111 |  |  |
|  |  | Interaction | ΔPuberty | -0.015 | 0.006 | -2.42 | 0.015 |  |  |
|  |  | Interaction | ΔPuberty:SexMale | -0.013 | 0.009 | -1.49 | 0.136 |  |  |
| Cortical | r.V1↔ES | Base | Age_1 | -0.004 | 0.002 | -2.85 | 0.004 | 0.573 | 0.601 |
|  |  | Base | ΔAge | 0.014 | 0.005 | 2.88 | 0.004 |  |  |
|  |  | Puberty | Age_1 | -0.005 | 0.002 | -2.79 | 0.005 |  |  |
|  |  | Puberty | ΔAge | 0.013 | 0.005 | 2.69 | 0.007 |  |  |
|  |  | Puberty | Puberty_1 | -0.001 | 0.002 | -0.37 | 0.708 |  |  |
|  |  | Puberty | ΔPuberty | 0.002 | 0.002 | 0.87 | 0.385 |  |  |
|  |  | Interaction | Age_1 | -0.005 | 0.002 | -2.81 | 0.005 |  |  |
|  |  | Interaction | ΔAge | 0.013 | 0.005 | 2.70 | 0.007 |  |  |
|  |  | Interaction | Puberty_1 | -0.001 | 0.002 | -0.36 | 0.717 |  |  |
|  |  | Interaction | ΔPuberty | 0.001 | 0.003 | 0.29 | 0.772 |  |  |
|  |  | Interaction | ΔPuberty:SexMale | 0.002 | 0.004 | 0.52 | 0.601 |  |  |
|  | r.LGN↔V1 | Base | Age_1 | 0.002 | 0.004 | 0.41 | 0.679 | 0.002 | 0.406 |
|  |  | Base | ΔAge | 0.009 | 0.012 | 0.73 | 0.464 |  |  |
|  |  | Puberty | Age_1 | -0.003 | 0.004 | -0.68 | 0.495 |  |  |
|  |  | Puberty | ΔAge | 0.002 | 0.013 | 0.19 | 0.847 |  |  |
|  |  | Puberty | Puberty_1 | 0.004 | 0.006 | 0.65 | 0.518 |  |  |
|  |  | Puberty | ΔPuberty | 0.018 | 0.005 | 3.58 | 3.5e-04 |  |  |
|  |  | Interaction | Age_1 | -0.003 | 0.004 | -0.72 | 0.473 |  |  |
|  |  | Interaction | ΔAge | 0.003 | 0.013 | 0.21 | 0.835 |  |  |
|  |  | Interaction | Puberty_1 | 0.004 | 0.006 | 0.66 | 0.506 |  |  |
|  |  | Interaction | ΔPuberty | 0.014 | 0.007 | 2.08 | 0.037 |  |  |
|  |  | Interaction | ΔPuberty:SexMale | 0.008 | 0.009 | 0.83 | 0.406 |  |  |
|  | r.ES↔IT | Base | Age_1 | 0.003 | 0.002 | 1.92 | 0.054 | 0.228 | 0.451 |
|  |  | Base | ΔAge | 0.016 | 0.005 | 3.26 | 0.001 |  |  |
|  |  | Puberty | Age_1 | 0.002 | 0.002 | 1.20 | 0.230 |  |  |
|  |  | Puberty | ΔAge | 0.016 | 0.005 | 3.16 | 0.002 |  |  |
|  |  | Puberty | Puberty_1 | 0.004 | 0.003 | 1.64 | 0.101 |  |  |
|  |  | Puberty | ΔPuberty | 0.002 | 0.002 | 0.89 | 0.371 |  |  |
|  |  | Interaction | Age_1 | 0.002 | 0.002 | 1.23 | 0.218 |  |  |
|  |  | Interaction | ΔAge | 0.016 | 0.005 | 3.15 | 0.002 |  |  |
|  |  | Interaction | Puberty_1 | 0.004 | 0.003 | 1.62 | 0.105 |  |  |
|  |  | Interaction | ΔPuberty | 0.003 | 0.003 | 1.17 | 0.243 |  |  |
|  |  | Interaction | ΔPuberty:SexMale | -0.003 | 0.004 | -0.75 | 0.451 |  |  |
|  | r.Amyg↔IT | Base | Age_1 | -0.010 | 0.005 | -1.97 | 0.049 | 0.170 | 0.777 |
|  |  | Base | ΔAge | -0.017 | 0.017 | -0.99 | 0.322 |  |  |
|  |  | Puberty | Age_1 | -0.007 | 0.006 | -1.22 | 0.222 |  |  |
|  |  | Puberty | ΔAge | -0.016 | 0.017 | -0.95 | 0.343 |  |  |
|  |  | Puberty | Puberty_1 | -0.016 | 0.009 | -1.85 | 0.064 |  |  |
|  |  | Puberty | ΔPuberty | -0.005 | 0.007 | -0.77 | 0.442 |  |  |
|  |  | Interaction | Age_1 | -0.007 | 0.006 | -1.21 | 0.227 |  |  |
|  |  | Interaction | ΔAge | -0.016 | 0.017 | -0.95 | 0.340 |  |  |
|  |  | Interaction | Puberty_1 | -0.016 | 0.009 | -1.86 | 0.063 |  |  |
|  |  | Interaction | ΔPuberty | -0.003 | 0.009 | -0.38 | 0.706 |  |  |
|  |  | Interaction | ΔPuberty:SexMale | -0.004 | 0.013 | -0.28 | 0.777 |  |  |
|  | l.V1↔ES | Base | Age_1 | -0.003 | 0.002 | -1.64 | 0.100 | 0.229 | 0.834 |
|  |  | Base | ΔAge | 0.013 | 0.005 | 2.65 | 0.008 |  |  |
|  |  | Puberty | Age_1 | -0.003 | 0.002 | -2.00 | 0.045 |  |  |
|  |  | Puberty | ΔAge | 0.012 | 0.005 | 2.36 | 0.018 |  |  |
|  |  | Puberty | Puberty_1 | 0.000 | 0.003 | 0.18 | 0.859 |  |  |
|  |  | Puberty | ΔPuberty | 0.003 | 0.002 | 1.70 | 0.089 |  |  |
|  |  | Interaction | Age_1 | -0.003 | 0.002 | -1.99 | 0.047 |  |  |
|  |  | Interaction | ΔAge | 0.012 | 0.005 | 2.36 | 0.018 |  |  |
|  |  | Interaction | Puberty_1 | 0.000 | 0.003 | 0.17 | 0.863 |  |  |
|  |  | Interaction | ΔPuberty | 0.004 | 0.003 | 1.40 | 0.162 |  |  |
|  |  | Interaction | ΔPuberty:SexMale | -0.001 | 0.004 | -0.21 | 0.834 |  |  |
|  | l.LGN↔V1 | Base | Age_1 | 0.005 | 0.004 | 1.11 | 0.268 | 0.001 | 0.886 |
|  |  | Base | ΔAge | 0.028 | 0.013 | 2.12 | 0.034 |  |  |
|  |  | Puberty | Age_1 | -0.001 | 0.004 | -0.12 | 0.904 |  |  |
|  |  | Puberty | ΔAge | 0.021 | 0.013 | 1.57 | 0.116 |  |  |
|  |  | Puberty | Puberty_1 | 0.007 | 0.007 | 0.96 | 0.335 |  |  |
|  |  | Puberty | ΔPuberty | 0.019 | 0.005 | 3.62 | 2.9e-04 |  |  |
|  |  | Interaction | Age_1 | -0.001 | 0.004 | -0.11 | 0.909 |  |  |
|  |  | Interaction | ΔAge | 0.021 | 0.013 | 1.57 | 0.117 |  |  |
|  |  | Interaction | Puberty_1 | 0.006 | 0.007 | 0.96 | 0.337 |  |  |
|  |  | Interaction | ΔPuberty | 0.020 | 0.007 | 2.77 | 0.006 |  |  |
|  |  | Interaction | ΔPuberty:SexMale | -0.001 | 0.010 | -0.14 | 0.886 |  |  |
|  | l.ES↔IT | Base | Age_1 | 0.003 | 0.002 | 1.84 | 0.066 | 0.001 | 0.181 |
|  |  | Base | ΔAge | 0.014 | 0.005 | 2.73 | 0.006 |  |  |
|  |  | Puberty | Age_1 | 0.001 | 0.002 | 0.33 | 0.742 |  |  |
|  |  | Puberty | ΔAge | 0.012 | 0.005 | 2.34 | 0.019 |  |  |
|  |  | Puberty | Puberty_1 | 0.007 | 0.003 | 2.67 | 0.008 |  |  |
|  |  | Puberty | ΔPuberty | 0.006 | 0.002 | 3.10 | 0.002 |  |  |
|  |  | Interaction | Age_1 | 0.001 | 0.002 | 0.38 | 0.700 |  |  |
|  |  | Interaction | ΔAge | 0.012 | 0.005 | 2.32 | 0.021 |  |  |
|  |  | Interaction | Puberty_1 | 0.007 | 0.003 | 2.64 | 0.008 |  |  |
|  |  | Interaction | ΔPuberty | 0.009 | 0.003 | 3.19 | 0.001 |  |  |
|  |  | Interaction | ΔPuberty:SexMale | -0.005 | 0.004 | -1.34 | 0.181 |  |  |
|  | l.Amyg↔IT | Base | Age_1 | -0.006 | 0.005 | -1.28 | 0.200 | 0.935 | 0.591 |
|  |  | Base | ΔAge | 0.006 | 0.016 | 0.36 | 0.722 |  |  |
|  |  | Puberty | Age_1 | -0.007 | 0.005 | -1.33 | 0.185 |  |  |
|  |  | Puberty | ΔAge | 0.005 | 0.017 | 0.32 | 0.749 |  |  |
|  |  | Puberty | Puberty_1 | 0.002 | 0.008 | 0.29 | 0.773 |  |  |
|  |  | Puberty | ΔPuberty | 0.002 | 0.007 | 0.29 | 0.773 |  |  |
|  |  | Interaction | Age_1 | -0.007 | 0.005 | -1.35 | 0.178 |  |  |
|  |  | Interaction | ΔAge | 0.005 | 0.017 | 0.33 | 0.741 |  |  |
|  |  | Interaction | Puberty_1 | 0.003 | 0.008 | 0.30 | 0.764 |  |  |
|  |  | Interaction | ΔPuberty | -0.001 | 0.009 | -0.15 | 0.881 |  |  |
|  |  | Interaction | ΔPuberty:SexMale | 0.007 | 0.012 | 0.54 | 0.591 |  |  |

***Supplementary Table 2:*** *Complete results of GLMs comparing data-driven and null tractography algorithms, including all estimated coefficients.*

| Tract | Coefficient | Effect size | SE | *z* value | *p* | *p*FDR |
| --- | --- | --- | --- | --- | --- | --- |
| r.SC↔Pul | (Intercept) | 9.84 | 0.022 | 439.0 | 0 | 0 |
|  | Algorithm | -1.56 | 0.016 | -97.4 | 0 | 0 |
|  | ScannerPhilips | 0.24 | 0.029 | 8.3 | 9.3e-17 | 3.0e-16 |
|  | ScannerSiemens | 0.21 | 0.020 | 10.4 | 4.1e-25 | 1.4e-24 |
|  | SexMale | 0.08 | 0.016 | 5.1 | 3.3e-07 | 7.9e-07 |
|  | EthnicityBlack | -0.08 | 0.026 | -3.3 | 0.001 | 0.002 |
|  | EthnicityHispanic | -0.06 | 0.021 | -2.8 | 0.005 | 0.009 |
|  | EthnicityAsian | -0.05 | 0.065 | -0.8 | 0.451 | 0.485 |
|  | EthnicityOther | -0.05 | 0.028 | -1.9 | 0.059 | 0.080 |
| r.SC↔LGN | (Intercept) | 8.10 | 0.027 | 299.0 | 0 | 0 |
|  | Algorithm | 0.26 | 0.019 | 13.5 | 1.8e-41 | 6.7e-41 |
|  | ScannerPhilips | -0.07 | 0.035 | -1.9 | 0.060 | 0.080 |
|  | ScannerSiemens | -0.05 | 0.024 | -2.0 | 0.042 | 0.058 |
|  | SexMale | -0.05 | 0.019 | -2.7 | 0.007 | 0.011 |
|  | EthnicityBlack | 0.04 | 0.031 | 1.2 | 0.223 | 0.254 |
|  | EthnicityHispanic | 0.18 | 0.025 | 7.2 | 5.9e-13 | 1.7e-12 |
|  | EthnicityAsian | 0.14 | 0.078 | 1.8 | 0.068 | 0.089 |
|  | EthnicityOther | 0.05 | 0.033 | 1.6 | 0.116 | 0.149 |
| r.SC↔Amyg | (Intercept) | 7.08 | 0.027 | 260.8 | 0 | 0 |
|  | Algorithm | -1.71 | 0.019 | -88.5 | 0 | 0 |
|  | ScannerPhilips | -0.23 | 0.035 | -6.7 | 2.8e-11 | 7.8e-11 |
|  | ScannerSiemens | -0.26 | 0.024 | -10.9 | 1.2e-27 | 4.3e-27 |
|  | SexMale | -0.02 | 0.019 | -1.2 | 0.226 | 0.254 |
|  | EthnicityBlack | 0.00 | 0.031 | 0.2 | 0.874 | 0.874 |
|  | EthnicityHispanic | 0.18 | 0.025 | 7.2 | 4.4e-13 | 1.3e-12 |
|  | EthnicityAsian | 0.31 | 0.078 | 4.0 | 6.6e-05 | 1.3e-04 |
|  | EthnicityOther | 0.08 | 0.034 | 2.3 | 0.024 | 0.036 |
| r.Pul↔Amyg | (Intercept) | 6.11 | 0.014 | 441.3 | 0 | 0 |
|  | Algorithm | 2.82 | 0.010 | 285.3 | 0 | 0 |
|  | ScannerPhilips | -0.10 | 0.018 | -5.5 | 3.1e-08 | 7.7e-08 |
|  | ScannerSiemens | -0.25 | 0.012 | -20.2 | 5.0e-91 | 2.4e-90 |
|  | SexMale | -0.05 | 0.010 | -4.7 | 2.2e-06 | 5.0e-06 |
|  | EthnicityBlack | 0.06 | 0.016 | 4.0 | 5.6e-05 | 1.1e-04 |
|  | EthnicityHispanic | -0.01 | 0.013 | -0.6 | 0.530 | 0.553 |
|  | EthnicityAsian | -0.06 | 0.040 | -1.5 | 0.139 | 0.176 |
|  | EthnicityOther | -0.01 | 0.017 | -0.7 | 0.482 | 0.511 |
| l.SC↔Pul | (Intercept) | 9.96 | 0.022 | 462.3 | 0 | 0 |
|  | Algorithm | -1.57 | 0.015 | -102.5 | 0 | 0 |
|  | ScannerPhilips | 0.16 | 0.028 | 5.8 | 7.3e-09 | 1.9e-08 |
|  | ScannerSiemens | 0.06 | 0.019 | 3.0 | 0.002 | 0.004 |
|  | SexMale | 0.12 | 0.015 | 7.7 | 1.8e-14 | 5.7e-14 |
|  | EthnicityBlack | -0.10 | 0.025 | -3.9 | 8.9e-05 | 1.7e-04 |
|  | EthnicityHispanic | -0.02 | 0.020 | -0.8 | 0.400 | 0.436 |
|  | EthnicityAsian | 0.13 | 0.062 | 2.1 | 0.033 | 0.048 |
|  | EthnicityOther | -0.03 | 0.027 | -1.3 | 0.196 | 0.231 |
| l.SC↔LGN | (Intercept) | 8.12 | 0.023 | 348.6 | 0 | 0 |
|  | Algorithm | 0.25 | 0.017 | 14.8 | 1.9e-49 | 7.9e-49 |
|  | ScannerPhilips | -0.15 | 0.030 | -5.0 | 5.7e-07 | 1.3e-06 |
|  | ScannerSiemens | -0.05 | 0.021 | -2.5 | 0.014 | 0.021 |
|  | SexMale | -0.05 | 0.017 | -3.0 | 0.003 | 0.005 |
|  | EthnicityBlack | 0.04 | 0.027 | 1.4 | 0.167 | 0.207 |
|  | EthnicityHispanic | 0.09 | 0.022 | 4.3 | 1.6e-05 | 3.5e-05 |
|  | EthnicityAsian | 0.21 | 0.067 | 3.1 | 0.002 | 0.004 |
|  | EthnicityOther | 0.04 | 0.029 | 1.4 | 0.170 | 0.207 |
| l.SC↔Amyg | (Intercept) | 7.27 | 0.026 | 280.3 | 0 | 0 |
|  | Algorithm | -1.30 | 0.019 | -70.3 | 0 | 0 |
|  | ScannerPhilips | -0.20 | 0.034 | -5.9 | 2.9e-09 | 7.8e-09 |
|  | ScannerSiemens | -0.32 | 0.023 | -14.1 | 7.5e-45 | 3.0e-44 |
|  | SexMale | 0.00 | 0.019 | 0.2 | 0.808 | 0.819 |
|  | EthnicityBlack | -0.12 | 0.030 | -4.1 | 4.0e-05 | 8.4e-05 |
|  | EthnicityHispanic | 0.07 | 0.024 | 2.7 | 0.006 | 0.010 |
|  | EthnicityAsian | 0.08 | 0.075 | 1.0 | 0.296 | 0.328 |
|  | EthnicityOther | 0.08 | 0.032 | 2.4 | 0.017 | 0.026 |
| l.Pul↔Amyg | (Intercept) | 6.11 | 0.014 | 426.8 | 0 | 0 |
|  | Algorithm | 2.82 | 0.010 | 275.9 | 0 | 0 |
|  | ScannerPhilips | -0.05 | 0.019 | -2.7 | 0.007 | 0.011 |
|  | ScannerSiemens | -0.26 | 0.013 | -20.1 | 8.3e-90 | 3.8e-89 |
|  | SexMale | -0.01 | 0.010 | -1.3 | 0.187 | 0.224 |
|  | EthnicityBlack | 0.07 | 0.016 | 4.0 | 5.9e-05 | 1.2e-04 |
|  | EthnicityHispanic | 0.03 | 0.013 | 2.1 | 0.037 | 0.052 |
|  | EthnicityAsian | -0.05 | 0.041 | -1.3 | 0.208 | 0.241 |
|  | EthnicityOther | -0.01 | 0.018 | -0.5 | 0.622 | 0.640 |

***Supplementary Table 3:*** *Complete results of linear mixed models of the effect of tract structural connectivity on response times during the emotional visual N-Back task.*

| Pathway | Tract | Coefficient | Effect size | SE | df | *t* value | *p* | *p*FDR |
| --- | --- | --- | --- | --- | --- | --- | --- | --- |
| Subcortical | l.Pul↔Amyg | (Intercept) | 1046.099 | 31.622 | 17054.3 | 33.08 | 1.2e-232 | 3.0e-231 |
|  |  | log(FBC) | -19.684 | 10.134 | 17041.9 | -1.94 | 0.052 | 0.095 |
|  |  | TypeFace | -59.800 | 37.015 | 12709.9 | -1.62 | 0.106 | 0.182 |
|  |  | Timepoint2 | -217.298 | 39.777 | 13578.9 | -5.46 | 4.8e-08 | 2.0e-07 |
|  |  | SexMale | -13.160 | 3.305 | 4338.7 | -3.98 | 7.0e-05 | 2.0e-04 |
|  |  | EthnicityBlack | 33.635 | 5.325 | 4360.5 | 6.32 | 2.9e-10 | 2.1e-09 |
|  |  | EthnicityHispanic | 23.495 | 4.290 | 4335.9 | 5.48 | 4.6e-08 | 2.0e-07 |
|  |  | EthnicityAsian | -27.991 | 13.347 | 4383.8 | -2.10 | 0.036 | 0.076 |
|  |  | EthnicityOther | -0.891 | 5.679 | 4336.5 | -0.16 | 0.875 | 0.918 |
|  |  | log(FBC):TypeFace | -8.116 | 11.896 | 12709.9 | -0.68 | 0.495 | 0.663 |
|  |  | log(FBC):Timepoint2 | 33.769 | 12.797 | 13581.6 | 2.64 | 0.008 | 0.020 |
|  |  | TypeFace:Timepoint2 | 10.684 | 53.745 | 12709.9 | 0.20 | 0.842 | 0.918 |
|  |  | log(FBC):TypeFace:Timepoint2 | -1.363 | 17.288 | 12709.9 | -0.08 | 0.937 | 0.958 |
|  | r.Pul↔Amyg | (Intercept) | 1088.340 | 34.355 | 17025.3 | 31.68 | 4.7e-214 | 1.0e-212 |
|  |  | log(FBC) | -33.083 | 10.951 | 17005.7 | -3.02 | 0.003 | 0.006 |
|  |  | TypeFace | -103.047 | 40.337 | 12712.7 | -2.55 | 0.011 | 0.024 |
|  |  | Timepoint2 | -289.328 | 42.936 | 13599.0 | -6.74 | 1.7e-11 | 2.0e-10 |
|  |  | SexMale | -13.236 | 3.305 | 4340.3 | -4.00 | 6.3e-05 | 1.9e-04 |
|  |  | EthnicityBlack | 33.712 | 5.326 | 4362.1 | 6.33 | 2.7e-10 | 2.1e-09 |
|  |  | EthnicityHispanic | 23.389 | 4.293 | 4344.2 | 5.45 | 5.4e-08 | 2.1e-07 |
|  |  | EthnicityAsian | -27.447 | 13.348 | 4386.4 | -2.06 | 0.040 | 0.080 |
|  |  | EthnicityOther | -0.797 | 5.677 | 4334.7 | -0.14 | 0.888 | 0.924 |
|  |  | log(FBC):TypeFace | 5.775 | 12.893 | 12712.9 | 0.45 | 0.654 | 0.804 |
|  |  | log(FBC):Timepoint2 | 56.655 | 13.738 | 13601.9 | 4.12 | 3.7e-05 | 1.3e-04 |
|  |  | TypeFace:Timepoint2 | 54.143 | 57.904 | 12710.6 | 0.94 | 0.350 | 0.523 |
|  |  | log(FBC):TypeFace:Timepoint2 | -15.271 | 18.525 | 12710.6 | -0.82 | 0.410 | 0.592 |
| Cortical | l.Amyg↔IT | (Intercept) | 997.722 | 17.710 | 17061.4 | 56.34 | 0 | 0 |
|  |  | log(FBC) | -5.471 | 7.446 | 17029.1 | -0.73 | 0.463 | 0.633 |
|  |  | TypeFace | -75.833 | 20.714 | 12710.7 | -3.66 | 2.5e-04 | 6.6e-04 |
|  |  | Timepoint2 | -112.483 | 22.009 | 13566.6 | -5.11 | 3.3e-07 | 1.2e-06 |
|  |  | SexMale | -13.159 | 3.307 | 4338.7 | -3.98 | 7.0e-05 | 2.0e-04 |
|  |  | EthnicityBlack | 33.743 | 5.329 | 4362.0 | 6.33 | 2.7e-10 | 2.1e-09 |
|  |  | EthnicityHispanic | 23.749 | 4.292 | 4337.2 | 5.53 | 3.3e-08 | 1.7e-07 |
|  |  | EthnicityAsian | -27.490 | 13.342 | 4376.7 | -2.06 | 0.039 | 0.080 |
|  |  | EthnicityOther | -0.624 | 5.679 | 4331.9 | -0.11 | 0.913 | 0.942 |
|  |  | log(FBC):TypeFace | -3.905 | 8.763 | 12710.6 | -0.45 | 0.656 | 0.804 |
|  |  | log(FBC):Timepoint2 | 0.091 | 9.238 | 13575.0 | 0.01 | 0.992 | 0.992 |
|  |  | TypeFace:Timepoint2 | -8.425 | 29.745 | 12709.3 | -0.28 | 0.777 | 0.888 |
|  |  | log(FBC):TypeFace:Timepoint2 | 6.309 | 12.479 | 12709.3 | 0.51 | 0.613 | 0.781 |
|  | r.Amyg↔IT | (Intercept) | 994.369 | 16.634 | 17037.0 | 59.78 | 0 | 0 |
|  |  | log(FBC) | -3.847 | 6.848 | 17062.7 | -0.56 | 0.574 | 0.747 |
|  |  | TypeFace | -83.970 | 19.321 | 12710.8 | -4.35 | 1.4e-05 | 5.0e-05 |
|  |  | Timepoint2 | -135.953 | 20.533 | 13453.7 | -6.62 | 3.7e-11 | 4.0e-10 |
|  |  | SexMale | -13.413 | 3.309 | 4345.6 | -4.05 | 5.1e-05 | 1.6e-04 |
|  |  | EthnicityBlack | 33.527 | 5.329 | 4363.9 | 6.29 | 3.5e-10 | 2.3e-09 |
|  |  | EthnicityHispanic | 23.423 | 4.296 | 4344.4 | 5.45 | 5.2e-08 | 2.1e-07 |
|  |  | EthnicityAsian | -27.802 | 13.347 | 4380.5 | -2.08 | 0.037 | 0.077 |
|  |  | EthnicityOther | -0.956 | 5.680 | 4334.9 | -0.17 | 0.866 | 0.918 |
|  |  | log(FBC):TypeFace | -0.430 | 7.984 | 12710.6 | -0.05 | 0.957 | 0.964 |
|  |  | log(FBC):Timepoint2 | 9.633 | 8.390 | 13459.3 | 1.15 | 0.251 | 0.399 |
|  |  | TypeFace:Timepoint2 | -13.186 | 27.917 | 12710.4 | -0.47 | 0.637 | 0.796 |
|  |  | log(FBC):TypeFace:Timepoint2 | 8.009 | 11.401 | 12710.3 | 0.70 | 0.482 | 0.653 |
|  | l.LGN↔V1 | (Intercept) | 964.173 | 18.614 | 16363.8 | 51.80 | 0 | 0 |
|  |  | log(FBC) | 8.176 | 7.203 | 16539.1 | 1.14 | 0.256 | 0.402 |
|  |  | TypeFace | -61.128 | 20.951 | 12704.8 | -2.92 | 0.004 | 0.009 |
|  |  | Timepoint2 | -122.450 | 22.379 | 13215.5 | -5.47 | 4.5e-08 | 2.0e-07 |
|  |  | SexMale | -12.988 | 3.312 | 4342.6 | -3.92 | 8.9e-05 | 2.5e-04 |
|  |  | EthnicityBlack | 33.277 | 5.335 | 4364.4 | 6.24 | 4.9e-10 | 2.9e-09 |
|  |  | EthnicityHispanic | 23.059 | 4.308 | 4352.8 | 5.35 | 9.1e-08 | 3.5e-07 |
|  |  | EthnicityAsian | -28.391 | 13.361 | 4381.5 | -2.12 | 0.034 | 0.072 |
|  |  | EthnicityOther | -1.070 | 5.683 | 4331.6 | -0.19 | 0.851 | 0.918 |
|  |  | log(FBC):TypeFace | -9.360 | 8.164 | 12704.8 | -1.15 | 0.252 | 0.399 |
|  |  | log(FBC):Timepoint2 | 3.715 | 8.650 | 13207.4 | 0.43 | 0.668 | 0.811 |
|  |  | TypeFace:Timepoint2 | -12.984 | 30.817 | 12705.2 | -0.42 | 0.674 | 0.811 |
|  |  | log(FBC):TypeFace:Timepoint2 | 7.662 | 11.911 | 12705.2 | 0.64 | 0.520 | 0.683 |
|  | r.LGN↔V1 | (Intercept) | 980.555 | 18.644 | 15704.9 | 52.59 | 0 | 0 |
|  |  | log(FBC) | 1.727 | 7.231 | 15920.7 | 0.24 | 0.811 | 0.917 |
|  |  | TypeFace | -58.064 | 20.671 | 12708.3 | -2.81 | 0.005 | 0.012 |
|  |  | Timepoint2 | -80.211 | 21.557 | 13106.9 | -3.72 | 2.0e-04 | 5.3e-04 |
|  |  | SexMale | -13.678 | 3.317 | 4353.0 | -4.12 | 3.8e-05 | 1.3e-04 |
|  |  | EthnicityBlack | 34.076 | 5.337 | 4368.7 | 6.39 | 1.9e-10 | 1.6e-09 |
|  |  | EthnicityHispanic | 24.314 | 4.322 | 4370.5 | 5.63 | 2.0e-08 | 1.1e-07 |
|  |  | EthnicityAsian | -26.527 | 13.361 | 4383.7 | -1.99 | 0.047 | 0.089 |
|  |  | EthnicityOther | -0.402 | 5.684 | 4335.3 | -0.07 | 0.944 | 0.958 |
|  |  | log(FBC):TypeFace | -10.582 | 8.070 | 12708.4 | -1.31 | 0.190 | 0.308 |
|  |  | log(FBC):Timepoint2 | -12.582 | 8.380 | 13107.9 | -1.50 | 0.133 | 0.225 |
|  |  | TypeFace:Timepoint2 | -22.192 | 29.850 | 12708.4 | -0.74 | 0.457 | 0.633 |
|  |  | log(FBC):TypeFace:Timepoint2 | 11.260 | 11.603 | 12708.4 | 0.97 | 0.332 | 0.508 |
|  | l.V1↔ES | (Intercept) | 1234.681 | 80.150 | 15604.0 | 15.40 | 3.8e-53 | 5.4e-52 |
|  |  | log(FBC) | -52.582 | 16.866 | 15620.0 | -3.12 | 0.002 | 0.005 |
|  |  | TypeFace | -156.193 | 89.401 | 12710.3 | -1.75 | 0.081 | 0.146 |
|  |  | Timepoint2 | -215.713 | 94.980 | 13108.8 | -2.27 | 0.023 | 0.050 |
|  |  | SexMale | -13.454 | 3.301 | 4334.0 | -4.08 | 4.7e-05 | 1.5e-04 |
|  |  | EthnicityBlack | 34.068 | 5.322 | 4360.5 | 6.40 | 1.7e-10 | 1.6e-09 |
|  |  | EthnicityHispanic | 23.671 | 4.284 | 4331.3 | 5.53 | 3.5e-08 | 1.7e-07 |
|  |  | EthnicityAsian | -26.965 | 13.327 | 4376.7 | -2.02 | 0.043 | 0.082 |
|  |  | EthnicityOther | -1.074 | 5.671 | 4328.6 | -0.19 | 0.850 | 0.918 |
|  |  | log(FBC):TypeFace | 14.994 | 18.824 | 12710.3 | 0.80 | 0.426 | 0.602 |
|  |  | log(FBC):Timepoint2 | 22.093 | 19.874 | 13097.6 | 1.11 | 0.266 | 0.412 |
|  |  | TypeFace:Timepoint2 | 49.407 | 131.616 | 12708.9 | 0.38 | 0.707 | 0.844 |
|  |  | log(FBC):TypeFace:Timepoint2 | -9.107 | 27.546 | 12708.9 | -0.33 | 0.741 | 0.868 |
|  | r.V1↔ES | (Intercept) | 1353.070 | 85.351 | 15717.0 | 15.85 | 3.7e-56 | 6.8e-55 |
|  |  | log(FBC) | -77.005 | 17.843 | 15727.1 | -4.32 | 1.6e-05 | 5.6e-05 |
|  |  | TypeFace | -161.033 | 95.462 | 12709.8 | -1.69 | 0.092 | 0.159 |
|  |  | Timepoint2 | -253.592 | 100.195 | 13092.7 | -2.53 | 0.011 | 0.025 |
|  |  | SexMale | -13.370 | 3.299 | 4334.4 | -4.05 | 5.1e-05 | 1.6e-04 |
|  |  | EthnicityBlack | 34.449 | 5.320 | 4362.3 | 6.48 | 1.1e-10 | 1.1e-09 |
|  |  | EthnicityHispanic | 24.097 | 4.283 | 4333.2 | 5.63 | 2.0e-08 | 1.1e-07 |
|  |  | EthnicityAsian | -27.256 | 13.317 | 4377.3 | -2.05 | 0.041 | 0.080 |
|  |  | EthnicityOther | -1.009 | 5.666 | 4329.1 | -0.18 | 0.859 | 0.918 |
|  |  | log(FBC):TypeFace | 15.900 | 19.959 | 12709.9 | 0.80 | 0.426 | 0.602 |
|  |  | log(FBC):Timepoint2 | 30.042 | 20.823 | 13084.6 | 1.44 | 0.149 | 0.249 |
|  |  | TypeFace:Timepoint2 | 47.009 | 138.912 | 12709.0 | 0.34 | 0.735 | 0.868 |
|  |  | log(FBC):TypeFace:Timepoint2 | -8.557 | 28.873 | 12709.0 | -0.30 | 0.767 | 0.888 |
|  | l.ES↔IT | (Intercept) | 1181.285 | 75.436 | 16407.3 | 15.66 | 7.1e-55 | 1.2e-53 |
|  |  | log(FBC) | -48.034 | 18.435 | 16405.5 | -2.61 | 0.009 | 0.021 |
|  |  | TypeFace | -148.401 | 85.432 | 12706.5 | -1.74 | 0.082 | 0.147 |
|  |  | Timepoint2 | -234.085 | 90.441 | 13196.3 | -2.59 | 0.010 | 0.022 |
|  |  | SexMale | -12.647 | 3.312 | 4351.9 | -3.82 | 1.4e-04 | 3.8e-04 |
|  |  | EthnicityBlack | 33.530 | 5.321 | 4359.1 | 6.30 | 3.3e-10 | 2.2e-09 |
|  |  | EthnicityHispanic | 23.758 | 4.285 | 4330.5 | 5.54 | 3.1e-08 | 1.6e-07 |
|  |  | EthnicityAsian | -26.987 | 13.329 | 4376.1 | -2.02 | 0.043 | 0.082 |
|  |  | EthnicityOther | -0.922 | 5.671 | 4327.6 | -0.16 | 0.871 | 0.918 |
|  |  | log(FBC):TypeFace | 15.486 | 20.862 | 12706.5 | 0.74 | 0.458 | 0.633 |
|  |  | log(FBC):Timepoint2 | 29.884 | 21.968 | 13190.5 | 1.36 | 0.174 | 0.286 |
|  |  | TypeFace:Timepoint2 | 119.629 | 124.678 | 12707.1 | 0.96 | 0.337 | 0.510 |
|  |  | log(FBC):TypeFace:Timepoint2 | -27.518 | 30.287 | 12707.0 | -0.91 | 0.364 | 0.537 |
|  | r.ES↔IT | (Intercept) | 1128.156 | 84.550 | 16428.2 | 13.34 | 2.1e-40 | 2.7e-39 |
|  |  | log(FBC) | -34.276 | 20.203 | 16426.3 | -1.70 | 0.090 | 0.158 |
|  |  | TypeFace | -63.964 | 95.675 | 12703.8 | -0.67 | 0.504 | 0.668 |
|  |  | Timepoint2 | -83.120 | 99.593 | 13185.2 | -0.83 | 0.404 | 0.590 |
|  |  | SexMale | -12.507 | 3.315 | 4353.2 | -3.77 | 1.6e-04 | 4.4e-04 |
|  |  | EthnicityBlack | 33.284 | 5.321 | 4355.8 | 6.25 | 4.4e-10 | 2.7e-09 |
|  |  | EthnicityHispanic | 23.431 | 4.284 | 4326.1 | 5.47 | 4.8e-08 | 2.0e-07 |
|  |  | EthnicityAsian | -26.387 | 13.329 | 4373.6 | -1.98 | 0.048 | 0.089 |
|  |  | EthnicityOther | -0.904 | 5.669 | 4323.2 | -0.16 | 0.873 | 0.918 |
|  |  | log(FBC):TypeFace | -5.026 | 22.847 | 12703.8 | -0.22 | 0.826 | 0.918 |
|  |  | log(FBC):Timepoint2 | -6.644 | 23.672 | 13183.0 | -0.28 | 0.779 | 0.888 |
|  |  | TypeFace:Timepoint2 | -67.581 | 137.262 | 12702.9 | -0.49 | 0.622 | 0.786 |
|  |  | log(FBC):TypeFace:Timepoint2 | 17.580 | 32.627 | 12702.9 | 0.54 | 0.590 | 0.759 |

***Supplementary Table 4:*** *Results of model comparison between models with age and puberty terms and null models. For most tracts, the models with age and puberty terms explained significantly more variance in the change of structural connectivity than null models (with four fewer degrees of freedom, the number of added parameters).*

| Pathway | Tract | Sum of Squares Difference | *p* | *p*FDR |
| --- | --- | --- | --- | --- |
| Subcortical | r.Pul↔Amyg | 0.261 | 0.009 | 0.013 |
|  | l.Pul↔Amyg | 0.699 | 8.4e-07 | 8.4e-06 |
| Cortical | r.V1↔ES | 0.065 | 0.001 | 0.004 |
|  | r.LGN↔V1 | 0.328 | 0.009 | 0.013 |
|  | r.ES↔IT | 0.065 | 0.002 | 0.004 |
|  | r.Amyg↔IT | 0.377 | 0.078 | 0.087 |
|  | l.V1↔ES | 0.049 | 0.012 | 0.016 |
|  | l.LGN↔V1 | 0.518 | 8.5e-04 | 0.003 |
|  | l.ES↔IT | 0.103 | 6.8e-05 | 3.4e-04 |
|  | l.Amyg↔IT | 0.081 | 0.752 | 0.752 |

***Supplementary Table 5:*** *Complete results of models of the effects of baseline age, change in age (follow-up minus baseline), baseline pubertal development, and change in pubertal development, on change in tract structural connectivity, between baseline and follow-up.*

| Pathway | Tract | Coefficient | Effect size | SE | *t* value | *p* | *p*FDR |
| --- | --- | --- | --- | --- | --- | --- | --- |
| Subcortical | r.Pul↔Amyg | (Intercept) | 1.388 | 0.055 | 25.07 | 1.6e-129 | 1.2e-128 |
|  |  | log(FBC) | 0.571 | 0.012 | 47.83 | 0 | 0 |
|  |  | Age_1 | -0.003 | 0.004 | -0.76 | 0.447 | 0.546 |
|  |  | ΔAge | -0.003 | 0.011 | -0.24 | 0.809 | 0.858 |
|  |  | Puberty_1 | -0.006 | 0.006 | -1.10 | 0.272 | 0.386 |
|  |  | ΔPuberty | -0.013 | 0.004 | -3.05 | 0.002 | 0.007 |
|  |  | SexMale | 0.001 | 0.005 | 0.18 | 0.860 | 0.880 |
|  |  | EthnicityBlack | 0.008 | 0.007 | 1.16 | 0.245 | 0.366 |
|  |  | EthnicityHispanic | -0.013 | 0.006 | -2.37 | 0.018 | 0.042 |
|  |  | EthnicityAsian | -0.052 | 0.017 | -3.07 | 0.002 | 0.007 |
|  |  | EthnicityOther | -0.017 | 0.007 | -2.27 | 0.023 | 0.051 |
|  | l.Pul↔Amyg | (Intercept) | 1.385 | 0.056 | 24.74 | 2.1e-126 | 1.4e-125 |
|  |  | log(FBC) | 0.574 | 0.011 | 50.16 | 0 | 0 |
|  |  | Age_1 | -0.006 | 0.004 | -1.63 | 0.102 | 0.176 |
|  |  | ΔAge | 0.013 | 0.012 | 1.15 | 0.252 | 0.369 |
|  |  | Puberty_1 | -0.009 | 0.006 | -1.56 | 0.119 | 0.197 |
|  |  | ΔPuberty | -0.021 | 0.005 | -4.64 | 3.7e-06 | 1.8e-05 |
|  |  | SexMale | -0.005 | 0.005 | -1.04 | 0.297 | 0.415 |
|  |  | EthnicityBlack | 0.012 | 0.008 | 1.64 | 0.100 | 0.176 |
|  |  | EthnicityHispanic | -0.011 | 0.006 | -1.93 | 0.053 | 0.102 |
|  |  | EthnicityAsian | -0.024 | 0.018 | -1.36 | 0.173 | 0.268 |
|  |  | EthnicityOther | -0.023 | 0.008 | -3.10 | 0.002 | 0.006 |
| Cortical | r.V1↔ES | (Intercept) | 1.054 | 0.041 | 25.76 | 3.3e-136 | 2.7e-135 |
|  |  | log(FBC) | 0.795 | 0.008 | 96.85 | 0 | 0 |
|  |  | Age_1 | -0.005 | 0.002 | -2.79 | 0.005 | 0.015 |
|  |  | ΔAge | 0.013 | 0.005 | 2.69 | 0.007 | 0.020 |
|  |  | Puberty_1 | -0.001 | 0.002 | -0.37 | 0.708 | 0.779 |
|  |  | ΔPuberty | 0.002 | 0.002 | 0.87 | 0.385 | 0.486 |
|  |  | SexMale | -0.001 | 0.002 | -0.67 | 0.504 | 0.591 |
|  |  | EthnicityBlack | -0.001 | 0.003 | -0.34 | 0.735 | 0.796 |
|  |  | EthnicityHispanic | -0.002 | 0.002 | -0.89 | 0.372 | 0.482 |
|  |  | EthnicityAsian | -0.016 | 0.007 | -2.10 | 0.036 | 0.073 |
|  |  | EthnicityOther | -0.004 | 0.003 | -1.36 | 0.174 | 0.268 |
|  | r.LGN↔V1 | (Intercept) | 0.590 | 0.052 | 11.37 | 1.5e-29 | 8.3e-29 |
|  |  | log(FBC) | 0.781 | 0.009 | 91.86 | 0 | 0 |
|  |  | Age_1 | -0.003 | 0.004 | -0.68 | 0.495 | 0.589 |
|  |  | ΔAge | 0.002 | 0.013 | 0.19 | 0.847 | 0.880 |
|  |  | Puberty_1 | 0.004 | 0.006 | 0.65 | 0.518 | 0.600 |
|  |  | ΔPuberty | 0.018 | 0.005 | 3.58 | 3.5e-04 | 0.001 |
|  |  | SexMale | -0.022 | 0.006 | -3.98 | 7.1e-05 | 3.0e-04 |
|  |  | EthnicityBlack | 0.018 | 0.008 | 2.21 | 0.027 | 0.058 |
|  |  | EthnicityHispanic | 0.029 | 0.006 | 4.52 | 6.4e-06 | 2.8e-05 |
|  |  | EthnicityAsian | 0.019 | 0.019 | 0.99 | 0.320 | 0.440 |
|  |  | EthnicityOther | 0.009 | 0.008 | 1.11 | 0.266 | 0.383 |
|  | r.ES↔IT | (Intercept) | 0.983 | 0.042 | 23.57 | 1.3e-115 | 8.2e-115 |
|  |  | log(FBC) | 0.759 | 0.010 | 79.75 | 0 | 0 |
|  |  | Age_1 | 0.002 | 0.002 | 1.20 | 0.230 | 0.348 |
|  |  | ΔAge | 0.016 | 0.005 | 3.16 | 0.002 | 0.006 |
|  |  | Puberty_1 | 0.004 | 0.003 | 1.64 | 0.101 | 0.176 |
|  |  | ΔPuberty | 0.002 | 0.002 | 0.89 | 0.371 | 0.482 |
|  |  | SexMale | 0.007 | 0.002 | 2.92 | 0.004 | 0.010 |
|  |  | EthnicityBlack | 0.003 | 0.003 | 0.87 | 0.387 | 0.486 |
|  |  | EthnicityHispanic | 0.002 | 0.002 | 0.91 | 0.364 | 0.482 |
|  |  | EthnicityAsian | 0.012 | 0.008 | 1.55 | 0.122 | 0.199 |
|  |  | EthnicityOther | 0.008 | 0.003 | 2.62 | 0.009 | 0.023 |
|  | l.V1↔ES | (Intercept) | 1.073 | 0.039 | 27.19 | 2.7e-150 | 2.7e-149 |
|  |  | log(FBC) | 0.787 | 0.008 | 99.64 | 0 | 0 |
|  |  | Age_1 | -0.003 | 0.002 | -2.00 | 0.045 | 0.089 |
|  |  | ΔAge | 0.012 | 0.005 | 2.36 | 0.018 | 0.042 |
|  |  | Puberty_1 | 0.000 | 0.003 | 0.18 | 0.859 | 0.880 |
|  |  | ΔPuberty | 0.003 | 0.002 | 1.70 | 0.089 | 0.163 |
|  |  | SexMale | 0.002 | 0.002 | 0.69 | 0.491 | 0.589 |
|  |  | EthnicityBlack | -0.003 | 0.003 | -0.78 | 0.436 | 0.541 |
|  |  | EthnicityHispanic | -0.009 | 0.002 | -3.47 | 5.2e-04 | 0.002 |
|  |  | EthnicityAsian | -0.005 | 0.008 | -0.61 | 0.541 | 0.619 |
|  |  | EthnicityOther | 0.000 | 0.003 | -0.09 | 0.928 | 0.928 |
|  | l.LGN↔V1 | (Intercept) | 0.705 | 0.055 | 12.80 | 8.1e-37 | 4.8e-36 |
|  |  | log(FBC) | 0.716 | 0.009 | 78.45 | 0 | 0 |
|  |  | Age_1 | -0.001 | 0.004 | -0.12 | 0.904 | 0.914 |
|  |  | ΔAge | 0.021 | 0.013 | 1.57 | 0.116 | 0.197 |
|  |  | Puberty_1 | 0.007 | 0.007 | 0.96 | 0.335 | 0.454 |
|  |  | ΔPuberty | 0.019 | 0.005 | 3.62 | 2.9e-04 | 0.001 |
|  |  | SexMale | -0.014 | 0.006 | -2.40 | 0.017 | 0.041 |
|  |  | EthnicityBlack | 0.019 | 0.009 | 2.13 | 0.033 | 0.069 |
|  |  | EthnicityHispanic | 0.030 | 0.007 | 4.53 | 6.1e-06 | 2.8e-05 |
|  |  | EthnicityAsian | 0.053 | 0.020 | 2.61 | 0.009 | 0.023 |
|  |  | EthnicityOther | 0.005 | 0.009 | 0.55 | 0.585 | 0.660 |
|  | l.ES↔IT | (Intercept) | 1.076 | 0.041 | 26.40 | 2.0e-142 | 1.8e-141 |
|  |  | log(FBC) | 0.734 | 0.009 | 80.11 | 0 | 0 |
|  |  | Age_1 | 0.001 | 0.002 | 0.33 | 0.742 | 0.796 |
|  |  | ΔAge | 0.012 | 0.005 | 2.34 | 0.019 | 0.043 |
|  |  | Puberty_1 | 0.007 | 0.003 | 2.67 | 0.008 | 0.020 |
|  |  | ΔPuberty | 0.006 | 0.002 | 3.10 | 0.002 | 0.006 |
|  |  | SexMale | 0.011 | 0.002 | 4.80 | 1.6e-06 | 8.4e-06 |
|  |  | EthnicityBlack | 0.007 | 0.003 | 2.02 | 0.044 | 0.088 |
|  |  | EthnicityHispanic | 0.004 | 0.003 | 1.46 | 0.144 | 0.230 |
|  |  | EthnicityAsian | -0.003 | 0.008 | -0.38 | 0.705 | 0.779 |
|  |  | EthnicityOther | 0.006 | 0.003 | 1.87 | 0.061 | 0.114 |

***Supplementary Table 6:*** *Complete results of the two linear mixed models of the overall effects of age and puberty variables on the change in structural connectivity across tracts in each pathway, controlling for the differences in mean structural connectivity of each tract (as well as sex and ethnicity, similar to other models). Note that the mean of one tract in each pathway is included in the intercept and not separately estimated.*

| Pathway | Coefficient | Effect size | SE | df | *t* value | *p* | *p*FDR |
| --- | --- | --- | --- | --- | --- | --- | --- |
| Cortical | (Intercept) | 1.026 | 0.022 | 9480.0 | 47.67 | 0 | 0 |
|  | log(FBC_1) | 0.753 | 0.004 | 25542.0 | 208.18 | 0 | 0 |
|  | log(FBC_l.LGN↔V1) | -0.381 | 0.006 | 25771.8 | -63.57 | 0 | 0 |
|  | log(FBC_l.V1↔ES) | 0.175 | 0.003 | 25209.7 | 54.59 | 0 | 0 |
|  | log(FBC_r.ES↔IT) | 0.020 | 0.002 | 21686.0 | 8.96 | 3.5e-19 | 1.1e-18 |
|  | log(FBC_r.LGN↔V1) | -0.401 | 0.006 | 25771.3 | -66.73 | 0 | 0 |
|  | log(FBC_r.V1↔ES) | 0.183 | 0.003 | 25307.8 | 55.41 | 0 | 0 |
|  | Age | -0.001 | 0.001 | 4287.9 | -0.80 | 0.426 | 0.459 |
|  | ΔAge | 0.013 | 0.004 | 4275.5 | 2.98 | 0.003 | 0.005 |
|  | Puberty_1 | 0.004 | 0.002 | 4275.9 | 1.67 | 0.095 | 0.121 |
|  | ΔPuberty | 0.008 | 0.002 | 4275.4 | 4.88 | 1.1e-06 | 3.1e-06 |
|  | SexMale | -0.003 | 0.002 | 4275.7 | -1.67 | 0.095 | 0.121 |
|  | EthnicityBlack | 0.007 | 0.003 | 4278.7 | 2.59 | 0.010 | 0.015 |
|  | EthnicityHispanic | 0.009 | 0.002 | 4290.2 | 4.23 | 2.3e-05 | 5.5e-05 |
|  | EthnicityAsian | 0.010 | 0.007 | 4280.1 | 1.55 | 0.121 | 0.148 |
|  | EthnicityOther | 0.004 | 0.003 | 4276.5 | 1.39 | 0.165 | 0.192 |
| Subcortical | (Intercept) | 1.459 | 0.043 | 5664.4 | 33.65 | 1.7e-226 | 5.8e-226 |
|  | log(FBC_1) | 0.548 | 0.009 | 8279.1 | 64.37 | 0 | 0 |
|  | log(FBC_r.Pul↔Amyg) | 0.008 | 0.003 | 4269.6 | 3.11 | 0.002 | 0.004 |
|  | Age_1 | -0.004 | 0.003 | 4248.5 | -1.36 | 0.174 | 0.195 |
|  | ΔAge | 0.005 | 0.009 | 4245.2 | 0.49 | 0.625 | 0.625 |
|  | Puberty_1 | -0.008 | 0.005 | 4243.8 | -1.70 | 0.089 | 0.121 |
|  | ΔPuberty | -0.018 | 0.004 | 4243.6 | -4.80 | 1.7e-06 | 4.2e-06 |
|  | SexMale | -0.002 | 0.004 | 4243.5 | -0.49 | 0.622 | 0.625 |
|  | EthnicityBlack | 0.011 | 0.006 | 4243.5 | 1.78 | 0.075 | 0.110 |
|  | EthnicityHispanic | -0.013 | 0.005 | 4248.5 | -2.76 | 0.006 | 0.010 |
|  | EthnicityAsian | -0.040 | 0.014 | 4247.2 | -2.82 | 0.005 | 0.009 |
|  | EthnicityOther | -0.021 | 0.006 | 4245.1 | -3.39 | 6.9e-04 | 0.001 |
